# Paenilamicins from the honey bee pathogen *Paenibacillus larvae* are context-specific translocation inhibitors of protein synthesis

**DOI:** 10.1101/2024.05.21.595107

**Authors:** Timm O. Koller, Max J. Berger, Martino Morici, Helge Paternoga, Timur Bulatov, Adriana Di Stasi, Tam Dang, Andi Mainz, Karoline Raulf, Caillan Crowe-McAuliffe, Marco Scocchi, Mario Mardirossian, Bertrand Beckert, Nora Vázquez-Laslop, Alexander Mankin, Roderich D. Süssmuth, Daniel N. Wilson

## Abstract

The paenilamicins are a group of hybrid non-ribosomal peptide-polyketide compounds produced by the honey bee pathogen *Paenibacillus larvae* that display activity against Gram-positive pathogens, such as *Staphylococcus aureus*. While paenilamicins have been shown to inhibit protein synthesis, their mechanism of action has remained unclear. Here, we have determined structures of the paenilamicin PamB2 stalled ribosomes, revealing a unique binding site on the small 30S subunit located between the A- and P-site tRNAs. In addition to providing a precise description of interactions of PamB2 with the ribosome, the structures also rationalize the resistance mechanisms utilized by *P. larvae*. We could further demonstrate that PamB2 interferes with the translocation of mRNA and tRNAs through the ribosome during translation elongation, and that this inhibitory activity is influenced by the presence of modifications at position 37 of the A-site tRNA. Collectively, our study defines the paenilamicins as a new class of context-specific translocation inhibitors.

## Introduction

The increase in multi-drug resistance is making our current arsenal of clinically-relevant antibiotics obsolete^1^, with a significant contribution coming from so-called *ESKAPE* pathogens^2^. This problem is compounded by the rapid decline in the approval of new antibiotics, particularly those with novel scaffolds^1^, highlighting the need for the discovery and development of further antimicrobials. One potential class are the paenilamicins, a group of hybrid non-ribosomal peptide-polyketide compounds that display activity against Gram-positive bacteria, such as *Bacillus subtilis* and *Staphylococcus aureus*^3,4^. Paenilamicins were shown to be 4-fold more active against methicillin-resistant *S. aureus* than the gold-standard ciprofloxacin^4^, and also display antifungal activity against the opportunistic fungal pathogens *Sporobolomyces salmonicolor* and *Aspergillus fumigatus*^4^. Paenilamicins are produced by the bacterium *Paenibacillus larvae*, which is the causative agent of American Foulbrood – the most destructive bacterial brood disease affecting honey bees world-wide^5^. Infection assays using bee larvae and the insect pathogen *Bacillus thuringiensis* demonstrated that paenilamicin production by *P. larvae* is used for suppression of bacterial competitors during host infection^4^.

To date, four distinct paenilamicins have been structurally elucidated, PamA1, PamA2, PamB1 and PamB2, which comprise a common core built from 2,3,5-trihydroxy pentanoic acid (Hpa), D-alanine (D-Ala), *N*-methyl-diaminopropionic acid (mDap), galantinic acid (Gla), glycine (Gly) and 4,3-spermidine (Spd) (**Fig. 1a** and **Extended Data Fig. 1a**)^3,4^. In addition, PamB2 contains an N-terminal D-agmatine (D-Agm) and a central D-ornithine (D-Orn) (**Fig. 1a**), whereas the D-Orn is replaced with a D-lysine (D-Lys) in PamB1**(Extended Data Fig. 1a**)^3^. The D-Agm of PamB1 and PamB2 is substituted with cadaverine in PamA1 and PamA2, respectively^3^ (**Extended Data Fig. 1a**). The structure of the paenilamicins is closely related to that of galantin I (**Extended Data Fig. 1b**), a compound originally isolated from a soil sample from New Guinea in the 1970’s^6^. A recent structural revision of PamB2 revealed that the (*6R)*-configuration of the terminal amino group in Agm is important for maximal activity and plays a role in self-resistance, being required for acetylation, and thereby inactivation, by the self-resistance factor PamZ^4,7^. The protection of the N-terminus during biosynthesis by attachment of an Acyl-D-Asn moiety is also a prominent prodrug resistance mechanism^8^.

**Fig. 1:**
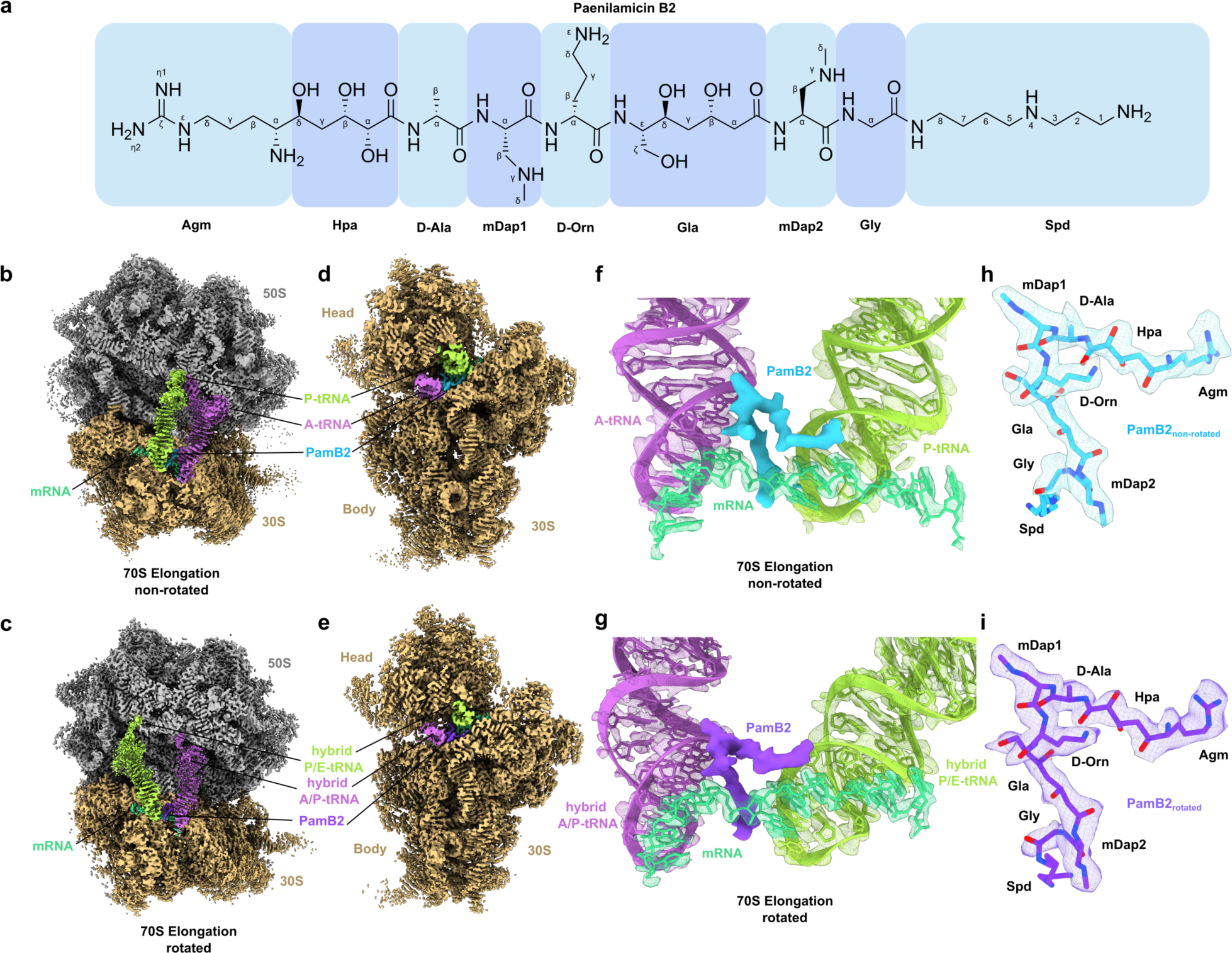
Cryo-EM structures of Paenilamicin B2 on the ribosome. **a**, Chemical structure of paenilamicin B2^3,4^. **b-d**, Transverse section of the cryo-EM map of the PamB2-stalled ribosome in non-rotated (**b,d**) and rotated (**c,e**) elongation state. **f-g**, Extracted cryo-EM density assigned to PamB2 from the non-rotated (**f**, light blue) and rotated (**g**, dark purple) with surrounding A-tRNA (purple), P-tRNA (light green) from the non-rotated (**f**) and hybrid A/P (purple) - and P/E-tRNA (light green) (**g**) and mRNA (cyan) in extracted density shown as mesh. **h**-**i**, Molecular model of PamB2 in extracted density of the non-rotated (**h**, light blue) and rotated (**i**, dark purple) states, shown as mesh.

The accordance of structural features of paenilamicins with the translation inhibitor edeine (**Extended Data Fig. 1c**) led to the hypothesis that paenilamicins may also exert their antimicrobial activity by binding to the ribosome and inhibiting protein synthesis^4^. In this regard, PamB2 was shown to be a potent inhibitor of *E. coli in vitro* translation systems, with a half-inhibitory concentration (IC_50_) of 0.4 μM^4^ – lower than reported for the well-known translation inhibitors erythromycin, chloramphenicol and tetracycline that display IC_50_ values of 0.75, 1.0, or 10 μM, respectively^9–11^. Although PamB2_2, which contains the non-native (*6S)*-configuration, retained inhibitory activity in the *E. coli in vitro* translation systems, a 7-10-fold reduction in efficiency (IC_50_ of 2.9 μM) was observed^4^. More dramatic was the effect of the PamZ-mediated acetylation of the 6-amino group of PamB2, which increased the IC_50_ by almost 80-fold to 31.9 μM^4^. This suggests that the acetylation of PamB2 may interfere with the binding of the compound to the ribosome. However, the binding site for PamB2 and the mechanism by which PamB2 inhibits protein synthesis remain to be elucidated.

Here, we have employed single particle cryo-EM to determine structures of ribosomes stalled during translation by the presence of PamB2 at 2.2-2.4 Å resolution. The structures reveal that PamB2 binds stably to elongation complexes that containing A- and P-site tRNAs, but not to initiating ribosomes bearing only a P-site tRNA, indicating that the presence of A-site tRNA is critical for the binding of PamB2 to the ribosome. This binding site of PamB2 is distinct from any other antibiotic binding site on the 30S subunit being located between the A- and P-site tRNAs. The structures also rationalize the increased activity of the native (*6R*)-configuration as well as the mechanism of self-resistance used by the producer. Together with complementary biochemical data, we demonstrate that PamB2 inhibits the EF-G catalyzed translocation step of protein synthesis in a highly context-specific manner that is dependent on the type of modifications that are present at position 37 of the A-site tRNA. To our knowledge, the paenilamicins represent the first class of context-specific translocation inhibitors that are influenced by the modification state of the tRNA.

## Results

### Cryo-EM structures of paenilamicin B2 on the ribosome

To investigate how paenilamicins inhibit translation, we generated PamB2-ribosome complexes for single particle cryo-EM analysis. Rather than forming complexes on vacant ribosomes, or with pre-defined functional states, we instead aimed to utilize more physiological complexes where translating ribosomes become stalled by the presence of PamB2. To achieve this, we performed *in vitro* translation reactions on *E. coli* ribosomes using the Met-Leu-Ile-Phe-stop-mRNA (MLIFstop-mRNA), a model template that we had previously used to successfully determine structures of drosocin-stalled ribosomal complexes^12^. Toeprinting was used to monitor the position of ribosomes on the MLIFstop-mRNA in the presence of increasing concentrations of synthetic PamB2 (**Extended Data Fig. 2**). As a positive control, we used thiostrepton that traps ribosomes on the AUG initiation codon^13–15^, and as a negative control, we included the inactive *N*-acetylated form of PamB2 (*N*-Ac-PamB2)^4^ (**Extended Data Fig. 2**). In the absence of drug, bands are evident for ribosomes on the AUG start codon, and the adjacent UUG (Leu) codon, suggesting that initiation and the first elongation step are slow on this mRNA or that the mRNA contains secondary structure in this region. In the presence of thiostrepton, a single strong band is observed that corresponds to ribosomes trapped on the AUG start codon (**Extended Data Fig. 2**), as expected^13–15^. By contrast, increasing concentrations of PamB2 led to a gradual loss of ribosomes at the AUG codon and an increase in ribosomes stalled one codon further with the UUG (Leu) codon in the P-site. This shift in ribosome positioning was not observed for the *N*-Ac-PamB2, where the pattern looks similar to the no-drug control, consistent with the inactivity of this compound^4^ (**Extended Data Fig. 2**). We also tested PamB2_2 with the non-native (*6S*)-configuration, which like *N*-Ac-PamB2, appeared to have little inhibitory activity in this assay (**Extended Data Fig. 2**).

Since there was little difference in the toeprinting at 25-100 μM PamB2, we formed PamB2-stalled ribosomal complexes (PamB2-SRCs) using a concentration of 50 μM for the inhibitor. After incubation, the reactions were centrifuged through sucrose cushions and the pelleted ribosomal complexes were subjected to single particle cryo-EM analysis. *In silico* sorting of the cryo-EM data revealed three main populations of ribosomal states, namely, non-rotated 70S ribosomes with P-site tRNA only (15 %), or with A- and P-site tRNAs (31 %), as well as a population containing rotated 70S ribosomes with A/P- and P/E-hybrid site tRNAs (17 %) (**Supplementary Fig. 1**), which after refinement yielded final reconstructions with average resolutions of 2.4 Å, 2.2 Å and 2.3 Å, respectively (**Fig. 1b-e, Extended Data Fig. 3a-I** and **Supplementary Table 1**). In both reconstructions containing two tRNAs, we observed additional density located between the A- and P-site tRNAs that could be unambiguously assigned to PamB2 (**Fig. 1f-i**). The density of PamB2 was well-resolved, enabling the orientation of the inhibitor to be determined, and the N-terminal Agm and Hpa as well as central D-Ala, D-Orn, mDap1 and mDap2 and Gla moieties to be modelled (**Fig. 1h-i** and **Supplementary Fig. 2**). The exception was the C-terminal Spd moiety that was poorly ordered in both maps, with density observed only at low thresholds (**Extended Data Fig. 3j-m**). No density for PamB2 was evident in the cryo-EM reconstruction where only one tRNA (the initiator tRNA in the P-site) was present, suggesting that PamB2 may require an A-site tRNA to bind stably to the ribosome.

### Interaction of PamB2 with the ribosomal P-site

The PamB2 binding site is located predominantly on the 30S subunit of the 70S ribosome, where it inserts into the cleft between the A- and P-site tRNAs (**Fig. 2a** and **Supplementary Fig. 2**). Although we describe the interactions of PamB2 for the non-rotated A- and P-site tRNA state, we note that within the limits of the resolution of the reconstructions, the binding mode of PamB2 is similar, if not identical, for the rotated A/P- and P/E-hybrid state (**Fig. 2b**). In both states, PamB2 is oriented with the Agm sidechain extending towards h24, while the central region of PamB2 runs parallel to the mRNA as well as one strand of nucleotides in h44 (**Fig. 2a**). The central mDap1 region of PamB2 interacts with H69 of the 23S rRNA, and then kinks such that the C-terminal (Gla-mDap2-Gly-Spd) region passes between the A- and P-site tRNAs, with the Spd moiety extending towards h31 (**Fig. 2a**). The kinked conformation of PamB2 is likely to be stabilized by three intramolecular hydrogen bonds (**Fig. 2c**), as well as two water-mediated interactions (**Fig. 2d**). The structural similarity with PamB2 (**Extended Data Fig. 1d-i**) suggests other paenilamicins (PamB1, PamA1 and PamA2) and also galantin I are likely to interact with the ribosome in the same manner.

**Fig. 2:**
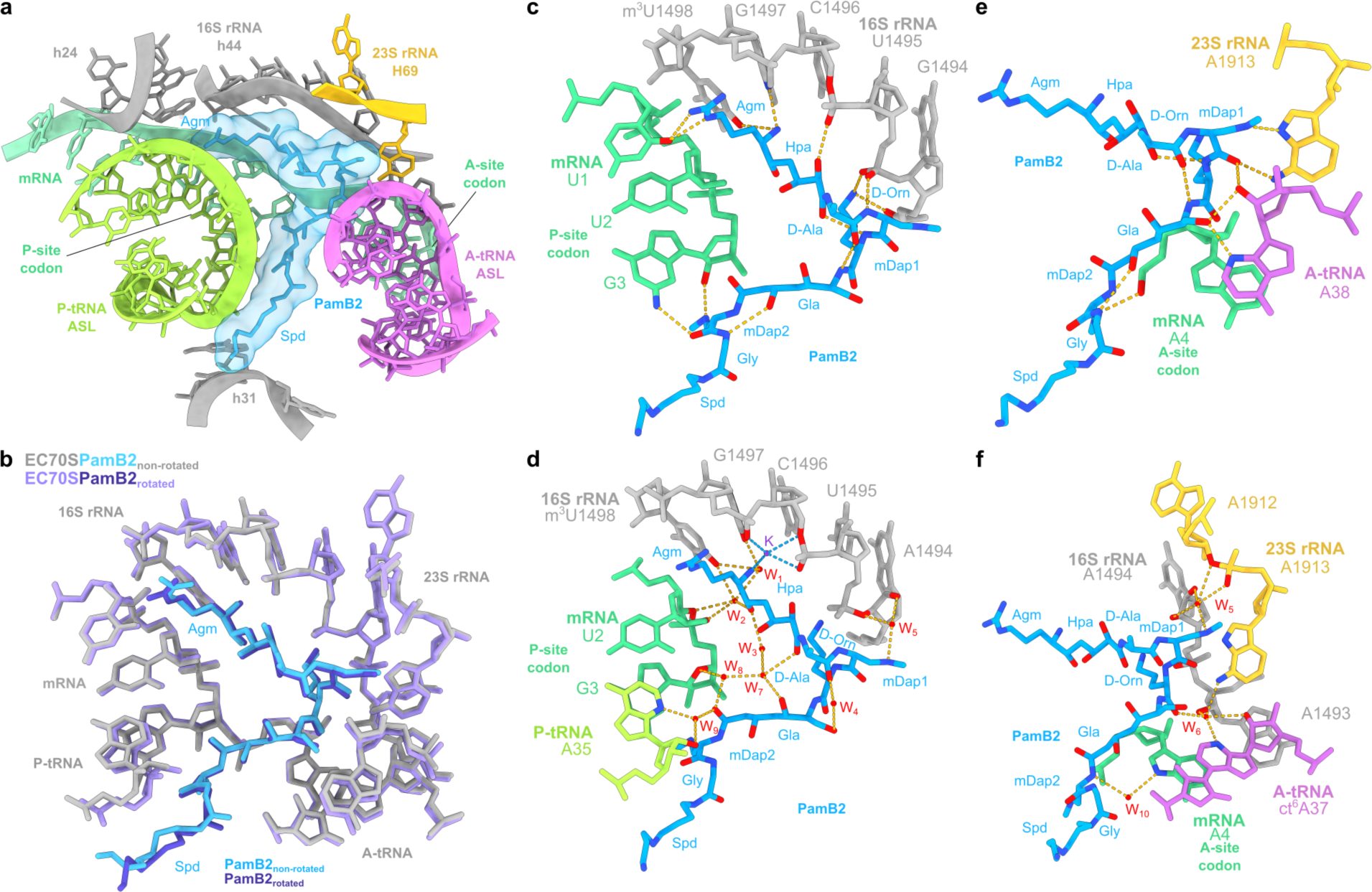
Interaction of PamB2 with the ribosomal P- and A-site. **a**, PamB2 (light blue) binding pocket located on the 30S subunit of the non-rotated PamB2-70S complex, with A-site tRNA (purple), P-site tRNA (light green), 16S rRNA (grey), 23S rRNA (yellow) and mRNA (cyan). **b**, Superimposition of the PamB2 binding pocket of the non-rotated (grey, with PamB2 in light blue) and rotated (light purple with PamB2 in purple) PamB2-70S complexes. **c-f**, Direct and water-mediated interactions (dashed yellow lines) between PamB2 and the ribosome, colored as in (**a**). **c**, Direct and intramolecular interactions of PamB2 with 16S rRNA of h44 and mRNA of the P-site codon. **d**, Water-mediated interactions of PamB2 with 16S rRNA of h44, mRNA of the P-site codon and P-site tRNA. **e**, Direct and intramolecular interactions of PamB2 with 23S rRNA of H69, mRNA of the A-site codon and A-site tRNA. **f**, Water-mediated interactions of PamB2 with 16S rRNA of h44, 23S rRNA of H69, mRNA of the A-site codon and A-site tRNA.

In the P-site, the majority of the interactions of PamB2 are with 16S rRNA nucleotides (G1494-m^3^U1498) in h44, on one side, and with the P-site codon of the mRNA, on the other (**Fig. 2c,d**). Together with U1495, C1496 and G1497, the N-terminal amino group of Agm coordinates an ion, which we assign to a K^+^ ion based on the coordination distances and the presence of a K^+^ ion in a similar position of a previous *E. coli* 70S-hygromycin B structure^16^. We note that acetylation of the N-terminal amino group of Agm by the *N*-acetyltransferase PamZ^4,7^, or modification with acyl-D-Asn^8^, inactivates PamB2. Modelling these modified forms of PamB2 into the binding site indicates that they would clash with the surrounding 16S rRNA (**Extended Data Fig. 4a-d**), suggesting that these modifications would prevent PamB2 from binding to the ribosome. The binding mode of PamB2 also explains the reduction in activity of PamB2_2 since the (*6S*)-configuration of the N-terminal amino group of Agm would lead to loss of direct contact with the N7 of G1497, as well as the K^+^ ion-mediated interaction with 16S rRNA nucleotides in h44 (**Extended Data Fig. 4e-f**).

With regard to the P-site codon of the mRNA, there are two main points of contact, namely, from the O2 of U1 (first position of the P-site codon) with the η2- and ε-nitrogens of Agm (**Fig. 2c**), and secondly, involving G3, located in the third position of the codon, where the ribose O2’ and N2 can form hydrogen bonds with γ-nitrogen and carbonyl-oxygen of mDap2 of PamB2 (**Fig. 2c**). In addition, water molecules (W_2_ and W_8_) mediate interactions between the backbone of U2 and Hpa of PamB2, as well as the O4’ (ribose) of G3 with the carbonyl-oxygen of Gla of PamB2 (**Fig. 2d**). Although PamB2 approaches the P-site tRNA, there is relatively little direct interaction, with the closest point of contact being 3.6 Å between the η2-nitrogen of Agm and the ribose 2’O of A37 of the P-site tRNA. However, we do observe a water-mediated (W_9_) interaction between the carbonyl-oxygen of Gla of PamB2 and the O2’ and N3 of A35 of the P-site tRNA (**Fig. 2d**).

### Interaction of PamB2 with the ribosomal A-site

In the A-site, PamB2 contacts not only with 16S rRNA nucleotides in h44, but also A1913 from H69 of the 23S rRNA, and, in contrast to the P-site, PamB2 makes extensive interactions with the A-site tRNA, albeit less with the mRNA codon (**Fig. 2c-f**). Interactions of PamB2 with the A-site tRNA revolve around nucleotides ct^6^A37 and A38, which are located in the anticodon-stem loop, directly adjacent to the anticodon (_34_CAU_36_) (**Fig. 2e,f**). Specifically, three direct hydrogen bonds are possible with A38 (**Fig. 2e**). Interactions with ct^6^A37 are indirect, being mediated by water W_6_, which is coordinated by the carbonyl-oxygen of D-Orn as well as the O2’ and N3 of ct^6^A37 (**Fig. 2f**). Interaction of PamB2 with the A-site codon of the mRNA is restricted to a direct interaction of backbone amide of Gly and a backbone oxygen of A4, which is located in the first position of the A-site codon, and a water-mediated interaction from the backbone amide of mDap2 via water W_10_ with N7 (3.4 Å) of A4 (**Fig. 2f**). The interactions between PamB2 and the A-site tRNA are likely to be critical for binding of PamB2 to the ribosome, since we observe no density for PamB2 in the P-tRNA-only state. We note that in structures of 70S ribosomes lacking A-tRNA^17^, the conformation of A1913 in H69 differs from that when the A-site tRNA is present, such that it would be incompatible with the interactions observed for PamB2 on the elongating ribosome (**Extended Data Fig. 4g-j**). The A1913 conformational shift induced by A-tRNA binding may therefore contribute to preventing stable binding of PamB2. Although a shift in the position of A1913 occurs during decoding when the A-site tRNA is still bound to EF-Tu, the A-tRNA itself is still sub-optimally placed to interact with PamB2 in such a state^18^ (**Extended Data Fig. 4k-l**), suggesting that full accommodation of A-tRNA is required for stable interaction of PamB2 with the ribosome.

### PamB2 inhibits tRNA_2_-mRNA translocation

Careful examination of the tRNAs in the PamB2-bound elongation complexes revealed the presence of additional density attached to the CCA-end of the A-site tRNA in the non-rotated elongation state and to the A/P-tRNA in the rotated hybrid state, indicating that peptide bond formation has already occurred in these complexes (**Supplementary Fig. 3a,b**). This suggests that PamB2 does not interfere with the decoding and accommodation by the A-tRNA, nor peptide bond formation, and also allows the ribosome to oscillate between the canonical and hybrid pre-translocational states (**Fig. 3a**). During normal translation, these pre-translocational states would be subject to the action of elongation factor EF-G, which binds and translocates the tRNA_2_-mRNA complex into the P- and E-sites, forming a post-translocational state^19–21^. The accumulation of pre-translocational states in the presence of PamB2, as well as the absence of post-translocation states (**Fig. 1b-e** and **Extended Data Fig. 5a-c**), suggests that PamB2 is likely to interfere with the process of translocation. To directly assess this, we analyzed the effect of PamB2 on EF-G-dependent translocation using the toeprinting assay. Ribosome complexes were formed with tRNA^fMet^ in the P-site and *N*-AcPhe-tRNA^Phe^ in the A-site, with toeprinting revealing a band corresponding to the expected pre-translocation state (**Fig. 3b**). In the absence of PamB2, but presence of EF-G, the toeprinting band shifted by three nucleotides, indicating that the A- and P-site tRNAs were translocated to the P- and E-sites (**Fig. 3b**). Little to no shift in the toeprint band was observed when the same reactions were performed in the presence of the control antibiotic negamycin, as reported previously^22^. Similarly, no shift in the toeprint was observed in the presence of PamB2, suggesting that PamB2 also interferes with the process of translocation (**Fig. 3b**). Comparison with recent structures of EF-G-bound translocation intermediates provides a structural rationale for the PamB2-mediated translocation inhibition^19–21^. While the initial binding of EF-G to the ribosome may be possible in the presence of PamB2 (**Extended Data Fig. 5d-f**)^20,21^, the subsequent steps where EF-G accommodates and promotes a shift in the anticodon stem of the A/P-site tRNA would lead to clashes with PamB2 (**Fig. 3c-e** and **Extended Data Fig. 5g-i**) ^19–21^. Moreover, in the early translocation intermediate with EF-G, A1913 rotates away from its position in the hybrid states^19–21^, which would require disruption of interactions between A1913 and PamB2 (**Extended Data Fig. 5j-l**). Collectively, this leads us to suggest that the interactions of PamB2 with the anticodon-stem loop of the A-site tRNA, as well as with the extended conformation of A1913, would prevent stable binding of EF-G to the pre-translational states, and thereby inhibit protein synthesis.

**Fig. 3:**
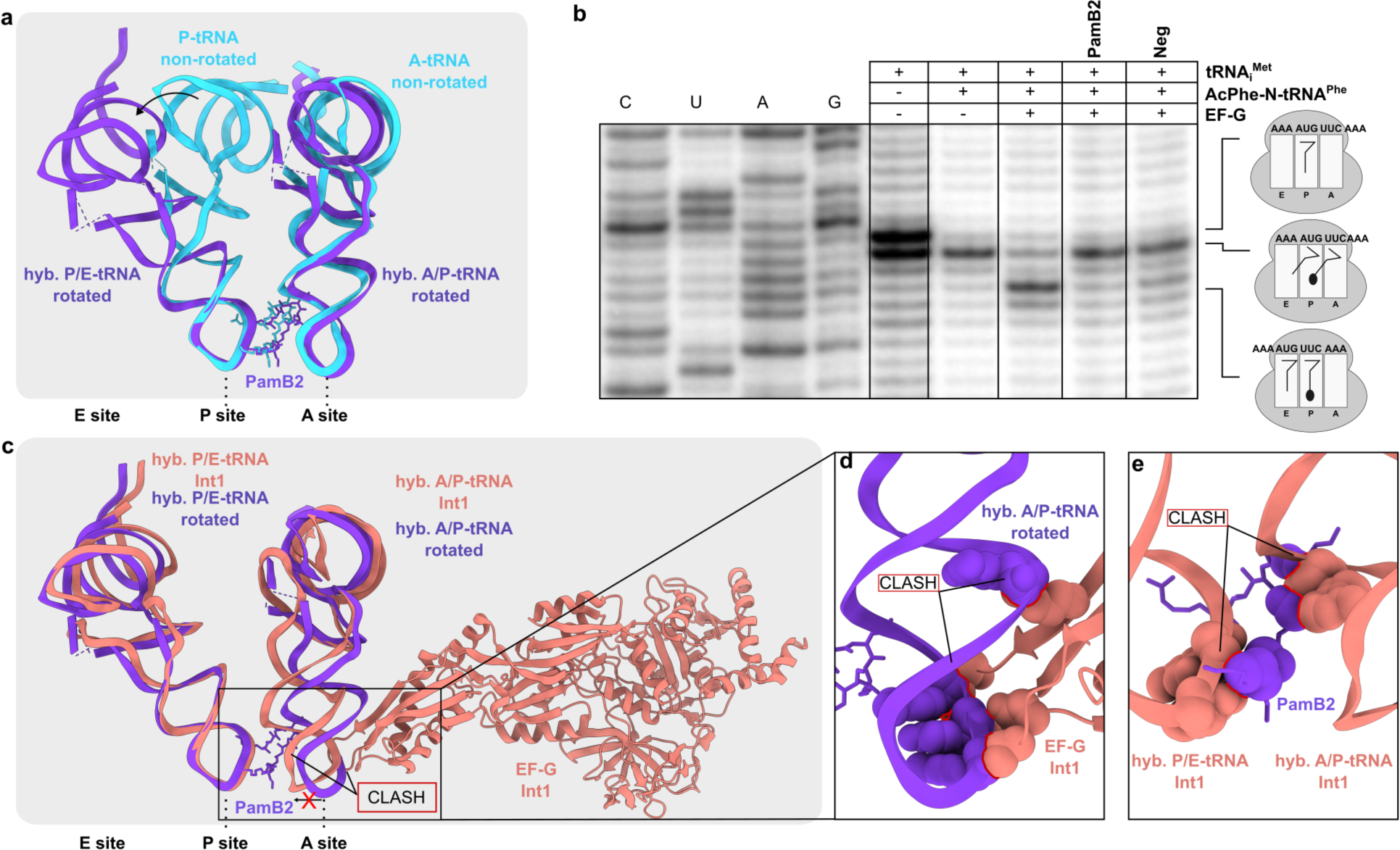
PamB2 inhibits tRNA_2_-mRNA translocation. **a**, Superimposition of PamB2 and tRNAs of the non-rotated (light blue) and rotated (purple) PamB2-70S complexes. **b**, Toeprinting assay monitoring the effect of PamB2 on EF-G dependent translocation, with initiator tRNA^fMet^ and N-AcPhe-tRNA^Phe^ in the absence of drugs and in the presence of the translocation inhibitor negamycin^22^. Toeprinting assays were performed in duplicate, with the duplicate gel present in the Source Data. **c**, Superimposition of PamB2 and hybrid tRNAs of the rotated (purple) PamB2-70S complex with hybrid tRNAs and EF-G bound to the *E. coli* 70S ribosome in the Int1 state (salmon, PDB ID 7N2V)^19^. **d,e**, Sphere representation of the (**d**) hybrid A/P-tRNA anticodon stem loop of the rotated PamB2 complex sterically clashing with EF-G (salmon, PDB ID 7N2V)^19^, and of the (**e**) hybrid A/P- and P/E-tRNAs of the Int1 state (PDB ID 7N2V)^19^ clashing with PamB2 (purple). Steric clashes are highlighted with red lines.

### Influence of A-site mRNA context on PamB2 inhibition

While the translocation assay (**Fig. 3b**) and structures of PamB2 bound to pre-translocation complexes (**Fig. 1b-e**) support the conclusion that PamB2 interferes with the EF-G mediated translocation process, our initial toeprinting assay indicated that it was not the first translocation step that was inhibited, but rather the second (**Extended Data Fig. 2**). If the first translocation reaction was inhibited, then ribosomes would be trapped with the AUG start codon in the P-site being decoded by the initiator tRNA^fMet^ and with a UUG codon in the A-site. While the density indicates that the initiator tRNA^fMet^ is present in the P-site of the P-tRNA-only volume (**Supplementary Fig. 4a**), the density for the mRNA codons and tRNAs in the A- and P-sites in structures of the PamB2-bound pre-translocation states indicated that one round of translocation had occurred before stalling of the complex (**Supplementary Fig. 4b-e**) i.e. UUG and AUA codons are in the P- and A-sites being decoded by tRNA^Leu^ (anticodon 5’-_35_CAA_37_-3’) and tRNA^Ile^ (anticodon 5’-_35_CAU_37_-3’), respectively (**Supplementary Fig. 4d-e**). Moreover, we observe extra density for the 2-methylthio-N6-isopentenyladenine (ms^2^i^6^A) at position 37 of the P-site tRNA^Leu^ as well as the cyclic N6-threonylcarbamoyladenosine (ct^6^A) at position 37 of the A-site tRNA^Ile^ (**Supplementary Fig. 4f-i**). Collectively, these findings suggest that PamB2 allowed the first translocation on the MLIFstop-mRNA, but prevented the second translocation reaction from taking place.

To investigate whether it is the initiation context that interferes with the action of PamB2, we generated a series of ErmBL mRNA templates containing 1-5 repeats of the UUG codon directly following the AUG start codon. In the absence of antibiotic, ribosomes initiate on the AUG start codon of the wildtype ErmBL mRNA (with one UUG repeat), and translate uninterrupted to the 12^th^ codon (AUC encoding Ile), where they become trapped due to the presence of the Ile-tRNA synthetase inhibitor mupirocin that was added to all reactions (**Fig. 4a**). As positive controls, we performed the assay with the wildtype ErmBL mRNA in the presence of the pleuromutilin retapamulin, which traps ribosomes on the AUG start codon^23^, as well as the macrolide erythromycin, which leads to the accumulation of ribosomes stalled with the 10^th^ CAU codon (encoding Asp) in the P-site (**Fig. 4a**), as we observed previously on the ErmBL mRNA^24,25^. Unlike retapamulin, the presence of PamB2 did not lead to a strong accumulation of ribosomes on the AUG start codon of the ErmBL mRNA, but rather ribosomes became stalled with the UUG codon in the P-site (**Fig. 4a**), as we observed for the MLIFstop-mRNA (**Extended Data Fig. 2**). While the insertion of UUG repeats into the ErmBL mRNA shifted the band for initiating ribosomes upwards in the gel as expected, the stalling bands remained constant (**Fig. 4a**), indicating that in the presence of PamB2, ribosomes can translate through stretches of up to five UUG codons unhindered. We conclude therefore that the lack of effect of PamB2 on the first translocation event in the wildtype ErmBL mRNA is not related to the initiation context, but apparently related to the presence of the UUG codon in the A-site. We also note that unlike for the short MLIFstop-mRNA, additional bands were observed on the longer ErmBL mRNA indicating that a subset of ribosomes also become stalled at subsequent sites in the mRNA, for example, with the 4^th^ UUC (encoding Phe) in the P-site, but not the 3^rd^ GUA codon in the P-site (**Fig. 4a**).

**Fig. 4:**
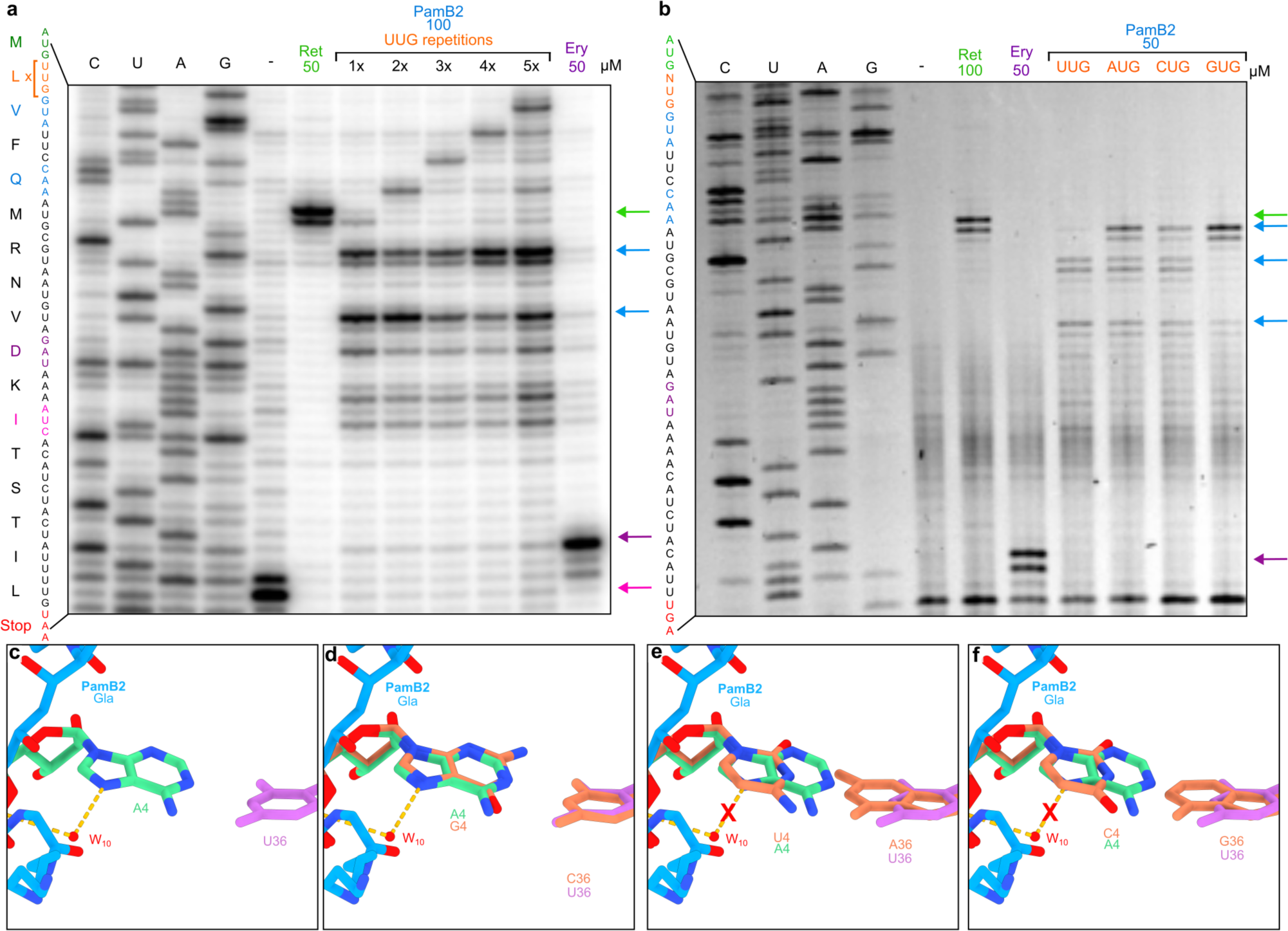
Influence of A-site mRNA context on PamB2 inhibition. **a-b**, Toeprinting assays monitoring the position of ribosomes on the wildtype ErmBL mRNA in the presence of water, 50 µM or 100 µM Retapamulin (Ret), 50 µM Erythromycin (Ery) and (**a**) an ErmBL mRNA with increasing number of UUG repetitions in the presence of 100 µM PamB2, and (**b**) with an ErmBL mRNA with the second codon mutated to UUG, AUG, CUG, GUG (orange) in the presence of 50 µM PamB2. Arrows indicate the stalling sites on the isoleucine catch codon in the presence of mupirocin (pink), at initiation (green), on the erythromycin-ErmBL stalling site (purple) and stalling induced by PamB2 (blue). The toeprinting assays were performed in duplicate. Toeprinting assays were performed in duplicate, with the duplicate gel present in the Source Data. **c-f,** Water (red) mediated interaction (dashed line) of PamB2 (blue) and the first nucleotide of the mRNA of the A-site codon (cyan), and superimposed with *in silico* mutated first position of the A-site codon (orange) to (**d**) guanosine, (**e**) uridine, or (**f**) cytidine. The loss of the water mediated interaction is indicated by a red cross.

An initial examination of the non-stalling contexts revealed that there are distinct sets of mRNA codons being recognized by distinct sets of tRNAs, namely, fMet-tRNA and Leu-tRNA decoding the AUG and UUG codons, respectively, in the first pre-translocational state, and Val-tRNA and Phe-tRNA decoding GUA and UUC codons, respectively, in the third pre-translocational state (**Fig. 4a**). Nevertheless, we noticed that in both contexts, the first position of the A-site codon was a uridine, whereas in the stalling contexts (UUG-GUA and UUC-CCA) using the ErmBL mRNA, the first position was either guanine or cytosine (**Fig. 4a**). In the PamB2-SRC that was generated on the MLIFstop-mRNA, the stalling context (UUG-AUA) had an adenine in the first position of the A-site codon (**Extended Data Fig. 1** and **Supplementary Fig. 4d-e**). Therefore, to test whether the nature of the A-site codon can influence the efficiency of PamB2-mediated translocation inhibition, we mutated the U in the first position of the A-site of the ErmBL mRNA to A, C and G (**Fig. 4b**). In contrast to U in the first position where little to no inhibition of the first translocation event was observed (**Fig. 4b**), clear toeprint bands were observed with each of the other nucleotides, indicating that ribosomes accumulate with the AUG start codon in the P-site when the A-site codon was changed from UUG to AUG, CUG or GUG (**Fig. 4b**). Although the inhibition by PamB2 with C in the first position appeared to be stronger than U, it was reproducibly weaker than with A and G (**Fig. 4b**). In fact, the inhibition with G in the first position of the A-site codon appeared to be almost complete, since no further stalling was observed at any of the downstream contexts (**Fig. 4b**). Although PamB2 does not directly interact with the first position of the A-site codon, we note that a water-mediated interaction is observed between the backbone amide of mDap2 via water W_10_ with the N7 of A in the first position of the A-site codon (**Fig. 4c**). Moreover, this interaction would be maintained with a G in the first position (**Fig. 4d**), where we observe strong inhibition (**Fig. 4b**), and would not be possible with U or C (**Fig. 4e-f**) where inhibition was weaker (**Fig. 4b**).

### Influence of A37 modification of A-tRNA on PamB2 inhibition

While the water-mediated interaction between PamB2 and the first position of the A-site codon may contribute to the specificity of stalling of PamB2, we note that it does not rationalize the difference in efficiency of inhibition of PamB2 that we observed between U and C in the first position (**Fig. 4b**). Therefore, we considered whether the nature of the tRNA in the A-site may also contribute, especially given that we observe interaction between PamB2 and nucleotides A37 and A38 of the A-site tRNA (**Fig. 2e, f**). Since we observe no inhibition by PamB2 when Phe-tRNA decodes UUC, we superimposed a ribosome structure with Phe-tRNA in the A-site^26^ and immediately noticed that tRNA^Phe^ bears a 2-methylthio-N6-isopentenyladenine (ms^2^i^6^A) at position 37, with the 2-methylthio moiety encroaching on the PamB2 binding site (**Fig. 5a-b**). In fact, with one exception (see later), all tRNAs that decode mRNA codons beginning with U have ms^2^i^6^A37, which is proposed to help stabilize the weaker U-C codon-anticodon interaction between the mRNA and the tRNA^27^. Consistently, we observe that tRNA^Leu^ that decodes UUG is also not inhibited by PamB2 (**Fig. 4a**) and would predict that similar results would be obtained for tRNA^Ser^ decoding UCU/UCA/UCG, tRNA^Tyr^ decoding UAU/UAC, tRNA^Cys^ decoding UGU/UGC and tRNA^Trp^ decoding UGG. The one exception is tRNA^Ser^ that decodes UCU and UCC where A37 is unmodified^27^. To directly test this, we generated a series of mRNA templates based on the ErmBL-(UUG)_4_ mRNA where we changed the 7^th^ GUA (Val) codon to each of the four serine codons UCC, UCU, UCA and UCG and performed the toeprinting assay in the presence of PamB2 (**Fig. 5c**). As hypothesized, strong stalling was observed at the UCC and UCU codons, which are decoded by the tRNA^Ser^ isoacceptor lacking any modification at position A37, whereas only weak stalling was observed at the UCA and UCG codons, which are decoded by the tRNA^Ser^ isoacceptor bearing ms^2^i^6^A37 (**Fig. 5c**). Thus, we conclude that PamB2 is a poor inhibitor of translocation when the A-site tRNA contains ms^2^i^6^A37 as a competitor.

**Fig. 5:**
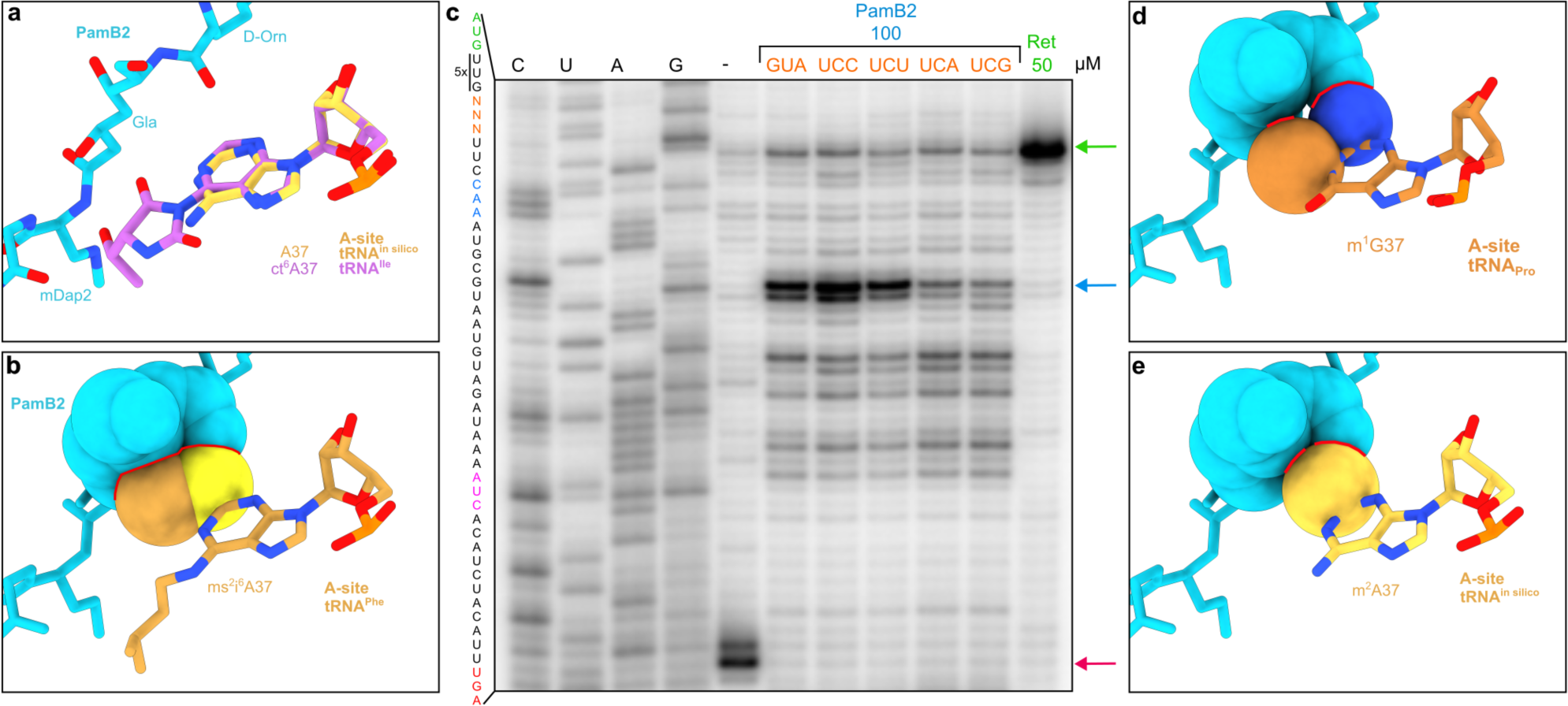
Influence of A37 modification of A-tRNA on PamB2 inhibition. **a**, PamB2 (light blue) and the modified A-site tRNA residue cyclic N6-threonylcarbamoyladenine (ct6) in position 37 (purple) from the non-rotated PamB2 complex superimposed with an *in silico* model of an unmodified A37. **b**, Superimposition of PamB2 from (**a**) with the 2-methylhio-N6-isopentenyladenine (ms^2^i^6^, light orange) at position 37 of the A-site tRNA^Phe^ from the *T. thermophilus* 70S ribosome pre-attack state (PDB ID 1VY5)^26^ shown as sphere representation with clashes indicated by red lines. **c**, Toeprinting assay monitoring the position of ribosomes on the (UUG)_5_-ErmBL mRNA in the presence of water, 50 µM retapamulin (Ret, green) and 100 µM PamB2 (light blue). The seventh codon was modified to different serine codons (orange). Arrows indicate stalling for the isoleucine catch codon in the presence of mupirocin (pink), the initiation (green) and PamB2-induced stalling (light blue). Toeprinting assays were performed in duplicate, with the duplicate gel present in the Source Data. **d-e**, Superimposition of PamB2 from (**a**) with (**d**) 1-methyl-guanine (m^1^G, dark orange) at position 37 of the A-site tRNA^Pro^ on the *T. thermophilus* 70S ribosome (PDB ID 6NUO)^40^ and (**e**) an *in silico* modified 2-methyl-adenine (m^2^A, yellow) shown as sphere representation with steric clashes indicated by red lines.

Although PamB2 was observed to inhibit translation when tRNA^Leu^ decoded CUG in the A-site, the extent of inhibition was relatively weak (**Fig. 4a**). Therefore, we superimposed a ribosome structure with tRNA^Leu^ in the A-site^17^ and recognized that tRNA^Leu^ contains a m^1^G at position 37 instead of an A, causing a steric clash between the N2 group of m^1^G37 of the A-tRNA and the D-Orn of PamB2 (**Fig. 5d**). In fact, most tRNAs decoding CNN codons, including CUN by tRNA^Leu^, CCN by tRNA^Pro^ as well as CGG by tRNA^Arg^ contain m^1^G37 (**Supplementary Fig. 5**)^27^. The exceptions are tRNA^His^ and tRNA^Gln^ that decode CAU/C and CAA/G, as well as tRNA^Arg^ that decodes CGU/C/A, however, all of these tRNAs have m^2^A37^27^ (**Supplementary Fig. 5**) that would also be predicted to clash with the D-Orn of PamB2, similar to m^1^G37 (**Fig. 5d,e** and **Supplementary Fig. 5**). Collectively, we conclude that the efficiency of translocation inhibition by PamB2 is directly influenced by the nature of the A-site tRNA and in particular by modifications at position A37, such as m^1^G37, m^2^A37, but especially ms^2^i^6^A37 (**Supplementary Fig. 5**), where the steric overlap with the drug is largest.

## Discussion

Based on our results, we propose a model for the mechanism of action of how PamB2 binds to the ribosome and inhibits protein synthesis (**Fig. 6a-e**). Our data suggest that PamB2 does not interfere with translation initiation (**Fig. 6a**), nor the initial EF-Tu-mediated decoding steps during translation elongation (**Fig. 6a,b**), but rather binds stably to the ribosome once the A-site tRNA becomes accommodated on the large 50S subunit (**Fig. 6c**). Our structural data indicate that PamB2 does not prevent peptide bond formation (**Fig. 6c**), nor the ribosome from adopting the rotated conformation with hybrid state A/P- and P/E-site tRNAs (**Fig. 6d**). Instead, we demonstrate that PamB2 interferes with the subsequent translocation step, where the tRNA_2_-mRNA complex is moved through the ribosome to occupy the P- and E-sites (**Fig. 6e**). We suggest that translocation is inhibited because PamB2 traps a pre-translocational state that is incompatible with stable binding of elongation factor EF-G (**Fig. 6d,e** and **Fig. 3c-e**). Based on the high similarity in chemical structures (**Extended Data Fig. 1**), we propose that the mechanism of action described here for PamB2 will be similar, if not identical, for other paenilamicin congeners (PamB1, PamA1 and PamA2)^3^ as well as the related compound galantin I^6^. Notably, modifications at the N-terminal amine of Agm/Cad in each of these congeners with either an acyl-D-Asn^8^ or an acetyl moiety^7^ will interfere with ribosome binding (**Extended Data Fig. 4a-d**), which thus rationalizes the self-resistance strategies of *P. larvae*.

**Fig. 6:**
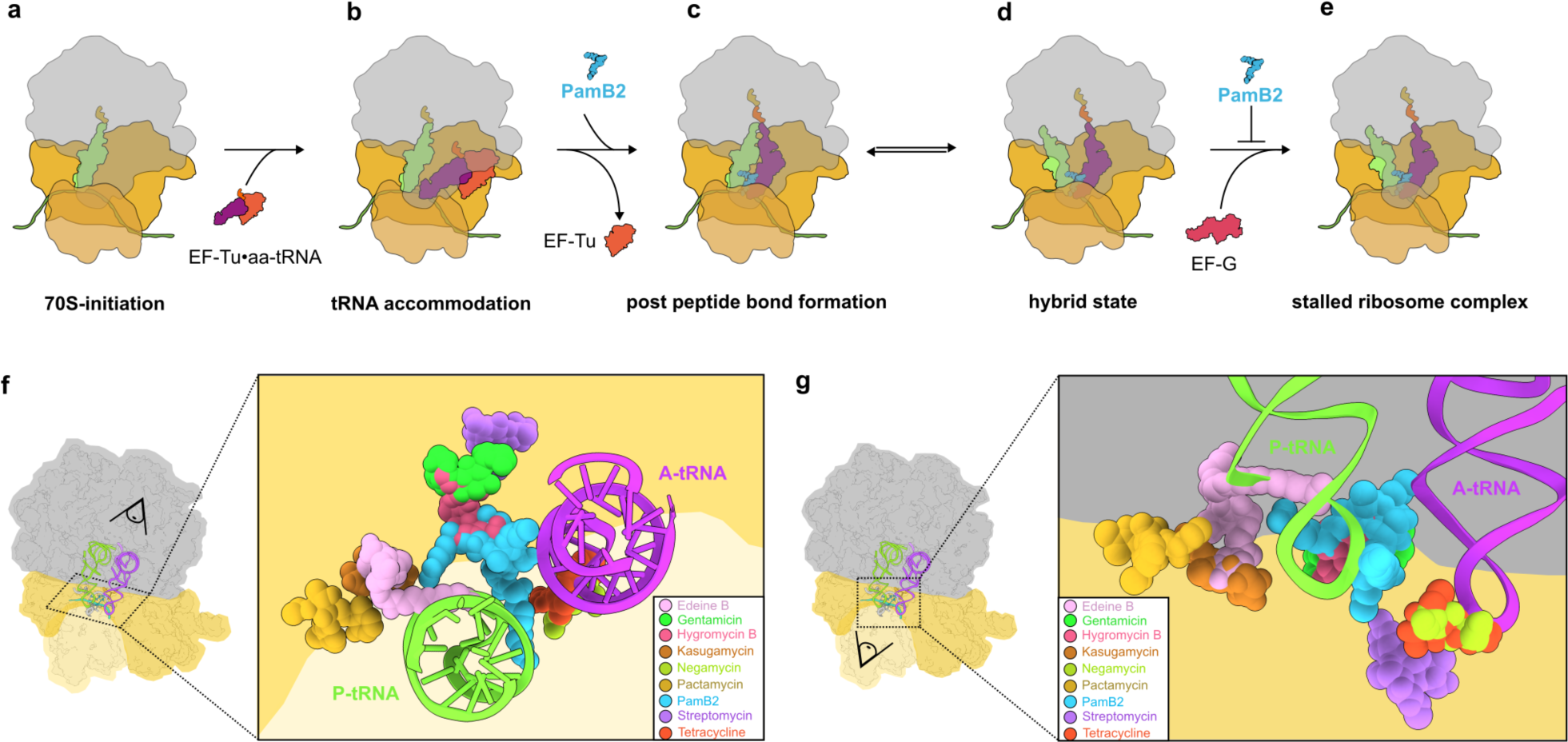
Mechanism of action of PamB2 and relative binding site of PamB2 compared to other antibiotics. **a-e**, Model for the mechanism of action of PamB2 during translation. **a-b**, PamB2 does not bind stably to (**a**) the initiation state with P-site tRNA only, nor (**b**) during delivery and decoding of the aminoacyl-tRNA to the A-site by EF-Tu. **c-d**, PamB2 binds stably to pre-translocation states with (**c**) A- and P-site tRNAs in non-rotated state and does not prevent peptide bond formation, as well as (**d**) rotated hybrid state with A/P- and P/E-tRNAs. **e**, Stable binding of EF-G is prevented by PamB2 thereby preventing translocation and trapping the ribosome in the pre-translocational states. **f-g**, Two views of PamB2 (light blue) superimposed with edeine B (pink, PDB ID 1I95)^41^, gentamicin (neon green, PDB ID 8CGU)^16^, hygromycin B (hot pink, PDB ID 8CAI)^16^, kasugamycin (dark orange, PDB ID 8CEP)^16^, negamycin (light green, PDB ID 4W2I)^22^, pactamycin (yellow, PDB ID 4W2H)^42^, streptomycin (pink, PDB ID 8CAI)^16^, and tetracycline (pink, PDB ID 8CF1)^16^, shown in sphere representation on the 30S subunit (head, light yellow; body, yellow), the 50S subunit (grey) and P-(green) and A-tRNA (purple).

One of the most unexpected findings of our study was that some pre-translocational states were refractory to the action of PamB2. Thus, PamB2 does not inhibit each and every round of translation elongation indiscriminately, but rather can be considered as a context-specific translocation inhibitor. The best understood context-specific translation inhibitors are those that target the large subunit, where their inhibitory action is influenced by the sequence of the nascent polypeptide chain being synthesized^28^, for example, macrolides and ketolides^29–31^, oxazolidinones and phenicols^32–34^ and more recently orthosomycins and tetracenomycins^35–37^. However, there are other examples of context-specific antibiotics that target the small subunit where their inhibitory activity appears to also be influenced by the nature of the mRNA and/or tRNA, such as pactamycin^38^, negamycin^22^ and kasugamycin^39^, yet, a structural basis for their specificity has so far been lacking. By contrast, we provide a structural basis for the context-specificity of PamB2, demonstrating that translation is less affected by the action of PamB2 when the A-site is occupied by tRNAs bearing modifications at the C2 position of nucleotide A37. This is exemplified by the majority of tRNAs decoding UNN codons that bear a ms^2^i^6^A37 modification, such that the 2-methylthio moiety would be predicted to sterically clash with the binding position of PamB2 on the ribosome (**Fig. 5a,b**). A weaker refractory action was also observed by tRNAs that decode CNN codons due to the presence of either m^1^G37 or m^2^A37, which would also lead to clashes with PamB2 (**Fig. 5d,e**). In *E. coli*, no other tRNAs have modifications at the A37 position that would be predicted to interfere with PamB2 activity^27^, however, we cannot exclude that it is different in other bacteria. Despite this context-specific action, PamB2 is a potent inhibitor of protein synthesis, with an IC_50_ of 0.4 μM, which is similar to, or better than, that reported for other well-known translation inhibitors, including erythromycin, chloramphenicol and tetracycline^4^. This is most likely because the translation of most, if not all, mRNAs involves translation of many codons that are read by tRNAs lacking modifications on the C2 position of A37 and are therefore susceptible to the action of PamB2.

Despite the suggested similarity to edeine^4^, we show here that the PamB2 binding site is completely unrelated to that reported for edeine (**Fig. 6f,g**). In fact, the binding site of PamB2 identified here is distinct from that reported for any other class of translation inhibitor (**Fig. 6f,g**). The only antibiotic with a binding site that slightly overlaps with PamB2 is the aminoglycoside hygromycin B (**Fig. 6f,g**), which binds within helix 44 and interacts with a putative K^+^ ion^16^. While this putative K^+^ ion is also coordinated by PamB2 (**Fig. 2d**), the majority of interactions with PamB2 are distinct from hygromycin B. The unique binding site and interactions of PamB2 with the ribosome compared to other clinically-used compounds, suggest that there is little chance for cross-resistance with PamB2. Together, with the good activity against methicillin-resistant *S. aureus*^4^, this makes paenilamicins an attractive class of compounds for the future development of novel antimicrobial agents to combat drug-resistant pathogenic bacteria.

## Acknowledgments

We thank A. Myasnikov, S. Nazarov and E. Ushikawa from the Dubochet Center for Imaging (an EPFL, UNIGE, UNIL initiative in Lausanne, Switzerland) for help with cryo-EM data collection and Dorota Klepacki for help with some of the toeprinting. This work was supported by the Deutsche Forschungsgemeinschaft (DFG) (grant WI3285/12-1 to D.N.W, grants SU239/21-1 and RTG 2473 “Bioactive Peptides” to R.D.S.) and the National Institute of Health (R35GM127134 to A.Man.). Part of this work was performed at the Multi-User CryoEM Facility at the Centre for Structural Systems Biology, Hamburg, supported by the Universität Hamburg and DFG grant numbers (INST152/772-1|152/774-1|152/775-1|152/776-1|152/777-1 FUGG).

## Author contributions

T.B., A.Mai. and R.S. synthesized PamB2 and T.D., A.Mai. and R.S. prepared the *N*-acetylated form of PamB2. M.J.B. prepared cryo-EM sample and performed toeprinting analysis. M.Mo., A.S., M.S., M.Ma, C.C-M. and K.R. performed activity studies. H.P. prepared cryo-EM grids and B.B. collected the cryo-EM data. T.O.K. processed the microscopy data, generated and refined the molecular models. T.O.K and M.J.B. prepared the figures. D.N.W. wrote the manuscript with input from all authors. D.N.W. and R.S. conceived and D.N.W., A.Man. and R.S. supervised the project.

## Additional information

### Supplementary information

The online version contains supplementary material available at https://…

## Methods

### Synthesis of Paenilamicins

Synthetic PamB and its *N*-acetylated form were produced as previously reported^4,7^

### Toeprinting assays

Toeprinting reactions for **Extended Data Fig. 2** and **Fig. 4b** were performed as described previously^14^. Briefly, reactions were performed with 6 μl of PURExpress *in vitro* protein synthesis system (New England Biolabs). The reactions were carried out on different templates (**Supplementary Table 2**). The reactions contained 340 ng of the respective mRNA template and were supplemented with the different compounds as specified. The translation reactions were incubated for 30 min at 37°C. The reverse transcription reaction was carried out using AMV RT and primer NV*1-Alexa 647 (5’-GGTTATAATGAATTTTGCTTATTAAC-3’). The translation reactions were incubated with the reverse transcriptase and the primer for 20 min at 37°C. mRNA degradation was carried out by the addition of 1 µl of 5 M NaOH. The reactions were neutralized with 0.7 µl of 25% HCl, and nucleotide removal was performed with the QIAquick Nucleotide Removal Kit (Qiagen). The samples were dried under vacuum for 2 hours at 60°C for subsequent gel electrophoresis. The 6% acrylamide gels were scanned on a Typhoon scanner (GE Healthcare).

Toeprinting reactions for **Fig. 4a** and **Fig. 5c** were performed as described previously^43^. Briefly, reactions were performed with 5 μl of PURExpress *in vitro* protein synthesis system. The reactions contained 0.1 pmol of the respective DNA template (**Supplementary Table 2**) and were supplemented with retapamulin, erythromycin, mupirocin or PamB2 as specified. The transcription-translation reactions were incubated 20 min at 37°C. Subsequently, reverse transcription was performed for 10 min at 37°C using AMV RT and the radiolabeled NV*1-primer. Reactions were stopped by the addition of 1 µl of 10 M NaOH and then neutralized with 0.8 µl of concentrated HCl. Subsequently, 200 µL of the stop buffer (0.3 M sodium acetate [pH 5.5], 5 mM EDTA and 0.5% SDS) was added and phenol extraction was performed. The obtained cDNA was precipitated in ethanol for subsequent gel electrophoresis.

### Translocation assay

Translocation assays were performed as previously described^22^ using the MFKAFK template^44^ (**Supplementary Table 2**). Reactions were prepared by incubating tight-coupled ribosomes (0.7 µM) with mRNA (0.5 µM) and tRNA ^Met.^ (1 µM) for 20 min at 37°C in the Pure System Buffer (5 mM Potassium phosphate [pH 7.3], 9 mM Mg(OAc)_2_, 95 mM potassium glutamate, 5 mM NH_4_Cl, 0.5 mM CaCl_2_, 1 mM spermidine, 8 mM putrescine, and 1 mM dithiothreitol) and for additional 10 min at 37°C with 2 μM of N-acetyl-Phe-tRNA^Phe^. At the time of N-acetyl-Phe-tRNA^Phe^ addition, the reactions were supplemented with PamB2 or negamycin as specified. The translocation reaction was initiated by addition of 1 µL EF-G/GTP mixture (1.2 µM/3.2 mM). After 5 min of incubation at 30°C, 2 µL reverse transcriptase/dNTPs mixture was added. The reactions were stopped after another 5 min at 30°C by addition of 200 µL of the stop buffer and subsequent phenol extraction. The obtained cDNA was precipitated in ethanol for subsequent gel electrophoresis.

### Preparation of complexes for structural analysis

PamB2-ribosome complexes were generated by *in vitro* transcription-translation reactions in PURExpress *in vitro* protein synthesis system (New England Biolabs) with the same reaction mix as described earlier in the *toeprinting assays*. Complex formation reactions were carried out on MLIF-stop toeprint DNA template in a 48 µL reaction in presence of 50 µM PamB2. The reaction was incubated for 15 min at 37°C. The reaction volume was then split: 42 µL were used for complex generation and 6 µL were used for toeprinting analysis. Ribosome complexes were isolated by centrifugation in 900 µL sucrose gradient buffer (containing 40% sucrose, 50 mM HEPES-KOH, pH 7.4, 100 mM KOAc, 25 mM Mg(OAc)_2_ and 6 mM 2-mercaptoethanol) for 3 hours at 4°C with 80,000 x g in a Optima™ Max-XP Tabletop Ultracentrifuge with a TLA 120.2 rotor. The pelleted complex was resuspended in Hico buffer (50 mM HEPES-KOH, pH 7.4, 100 mM KOAc, 25 mM Mg(OAc)_2_ supplemented with PamB2 at the same concentrations used in the *in vitro* translation reaction), then incubated for 15 min at 37°C.

### Preparation of cryo-EM grids and data collection

Sample volumes of 3.5 μl (8 OD_260_ per ml) were applied to grids (Quantifoil, Cu, 300 mesh, R3/3 with 3 nm carbon) which had been freshly glow-discharged using a GloQube (Quorum Technologies) in negative charge mode at 25 mA for 90 sec. Sample vitrification was performed using an ethane-propane mixture (37:63) in a Vitrobot Mark IV (Thermo Fisher Scientific), the chamber was set to 4°C and 100% relative humidity and blotting was done for 3 sec with no drain or wait time. Frozen cryo-EM grids were imaged on a TFS 300kV Titan Krios at the Dubochet Center for Imaging EPFL (Lausanne, Switzerland). Images were collected on Falcon IV direct detection camera in counting mode using the EPU and AFIS data collection scheme with a magnification of 96,000 x and a total dose of 60 electrons per square angstrom (e^−^/Å^2^) for each exposure, and defocus ranging from −0.4 to −0.9 microns. In total, 7,638 movies were produced in EER format at a pixel size of 0.8 Å/pixel.

### Single-particle reconstruction of PamB2-stalled ribosome complexes

RELION v4.0.1^45^ was used for processing, unless otherwise specified. For motion correction, RELION’s implementation of MotionCor2 with 4×4 patches and for initial CTF estimation CTFFIND v4.1.14^46^ was employed. From 7,638 micrographs, 611,189 particles were picked using crYOLO v1.8.04b47 with a general model^47^. 562,816 ribosome-like particles were selected after 2D classification and extracted at 3x decimated pixel size (2.4 Å/pixel) (**Supplementary Fig. 1b,c**). An initial 3D refinement was done using a *E. coli* 70S reference map (EMD-12573)^29^. Particles were 3D classified for 100 iterations and resulted in four classes of which a non-rotated 70S class with A-, P- and E-site tRNAs (65.0%, 365,773 particles) and a rotated 70S with hybrid A/P- and P/E-tRNA (22.4%, 126,259 particles) (**Supplementary Fig. 1d**) were further sub-sorted. Sub-sorting was done for 100 iterations for both classes individually (**Supplementary Fig. 1e,g**), yielding two classes of non-rotated 70S with A-, P- and sub-stoichiometric E-site tRNA (52.5%, 295,568 particles) and rotated 70S class with hybrid A/P- and P/E-tRNA (16.7%, 93,773 particles), respectively. Focus-sorting was performed with partial particle subtraction using a mask surrounding the tRNAs for the particles containing non-rotated 70S with A-, P- and sub-stoichiometric E-site tRNA and 3D classified for 100 iterations yielding six classes. Classes containing A-, P- and E-site tRNA (31.4%, 176,827 particles), as well as classes containing just P-site tRNA (15.1%, 84,771 particles) were combined and further processed (**Supplementary Fig. 1f**). All resulting classes were 3D refined (with a solvent mask), CTF refined (4^th^ order aberration, anisotropic magnification and per-particle defocus value estimation), Bayesian polished, again CTF refined and after a final 3D refinement yielded a final average resolution of 2.2 Å (at FSC_0.143_) for the post-processed masked reconstruction of the non-rotated 70S complex containing A-, P- and sub. E-site tRNA (**Supplementary Fig. 1h**), a final average resolution of 2.4 Å (at FSC_0.143_) for the post-processed masked reconstruction of the 70S complex containing P-tRNA (**Supplementary Fig. 1i**) and a final average resolution of 2.3 Å (at FSC_0.143_) for the post-processed masked reconstruction of the rotated 70S complex containing hybrid A/P-, and P/E-tRNAs (**Supplementary Fig. 1f**). To estimate local resolution values Bsoft^48^ was used on the half-maps of the final reconstructions (blocres -sampling 0.8 -maxres -boc 20 -cutoff 0.143 -verbose 1 -origin 0,0,0 -Mask half_map1 half_map 2) (**Extended Data Fig. 3d-m**).

### Molecular modelling of the PamB2-ribosome complexes

The molecular models of the 30S and 50S ribosomal subunits were based on the *E. coli* 70S ribosome (PDB ID 7K00)^49^. PamB2 and *in silico* modified versions of paenilamicins were generated and restraints created using aceDRG^50^ and modelled *de novo*. The non-rotated and rotated 70S complexes were assembled with tRNA^Leu^ and tRNA^Ile^ used from the drosocin-stalled 70S complexes (PDB ID 8AM9)^12^. The initiation complex was assembled with an initiator fMet-tRNA (PDB ID 1VY4)^26^ in the P-site. Modifications of rRNA nucleotides and tRNA^Leu^ and tRNA^Ile^ were generated using aceDRG^50^. Starting models were rigid body fitted using ChimeraX 1.6.1^51^ and modelled using Coot 0.9.8.92^52^ from the CCP4 software suite v8.0.017^53^. The sequence for the tRNAs were adjusted based the appropriate anticodons corresponding to the mRNA. Final refinements were done in REFMAC 5^54^ using Servalcat v0.4.28^55^. The molecular models were validated using Phenix comprehensive Cryo-EM validation in Phenix 1.20.1-4487^56^.

### Figures

UCSF ChimeraX 1.6.1^51^ was used to isolate density, align molecular models and visualize density images and structural superpositions. Figures were assembled with Inkscape (latest development release, regularly updated).

## Data availability

Micrographs have been deposited as uncorrected frames in the Electron Microscopy Public Image Archive (EMPIAR) with the accession codes EMPIAR-12080 [https://www.ebi.ac.uk/pdbe/emdb/empiar/entry/12080]. Cryo-EM maps have been deposited in the Electron Microscopy Data Bank (EMDB) with accession codes EMD-18950 [https://www.ebi.ac.uk/pdbe/entry/emdb/EMD-18950] (Non-rotated 70S PamB2 complex), EMD-19004 [https://www.ebi.ac.uk/pdbe/entry/emdb/EMD-19004] (Rotated 70S PamB2 complex), and EMD-50296 [https://www.ebi.ac.uk/pdbe/entry/emdb/EMD-50296] (Initiation 70S complex). Molecular models have been deposited in the Protein Data Bank with accession codes 8R6C [https://doi.org/10.2210/pdb8R6C/pdb] (Non-rotated 70S PamB2 complex), 8R8M [https://doi.org/10.2210/pdb8R8M/pdb] (Rotated 70S PamB2 complex), 9FBV [https://doi.org/10.2210/pdb9FBV/pdb] (Initiation 70S complex). Structures from prior studies were used in this work for comparison, alignments and for modelling and are available in the Protein Data Bank, with PDB ID 7K00, 8AM9, 1VY4, 1VY5, 6NUO, 7N2V, 7N2U, 7N1P, 1I95, 8CGU, 8CAI, 8CEP, 4W2I, 4W2H, 8CF1, 6WD2, 6WD8, 6WD0, 7SSL, 7SSD, 7PJY, 7PJW, 7PJV, 4V6Z, 4V8D. Source data are provided with this paper.

---

**Extended Data Fig. 1:**
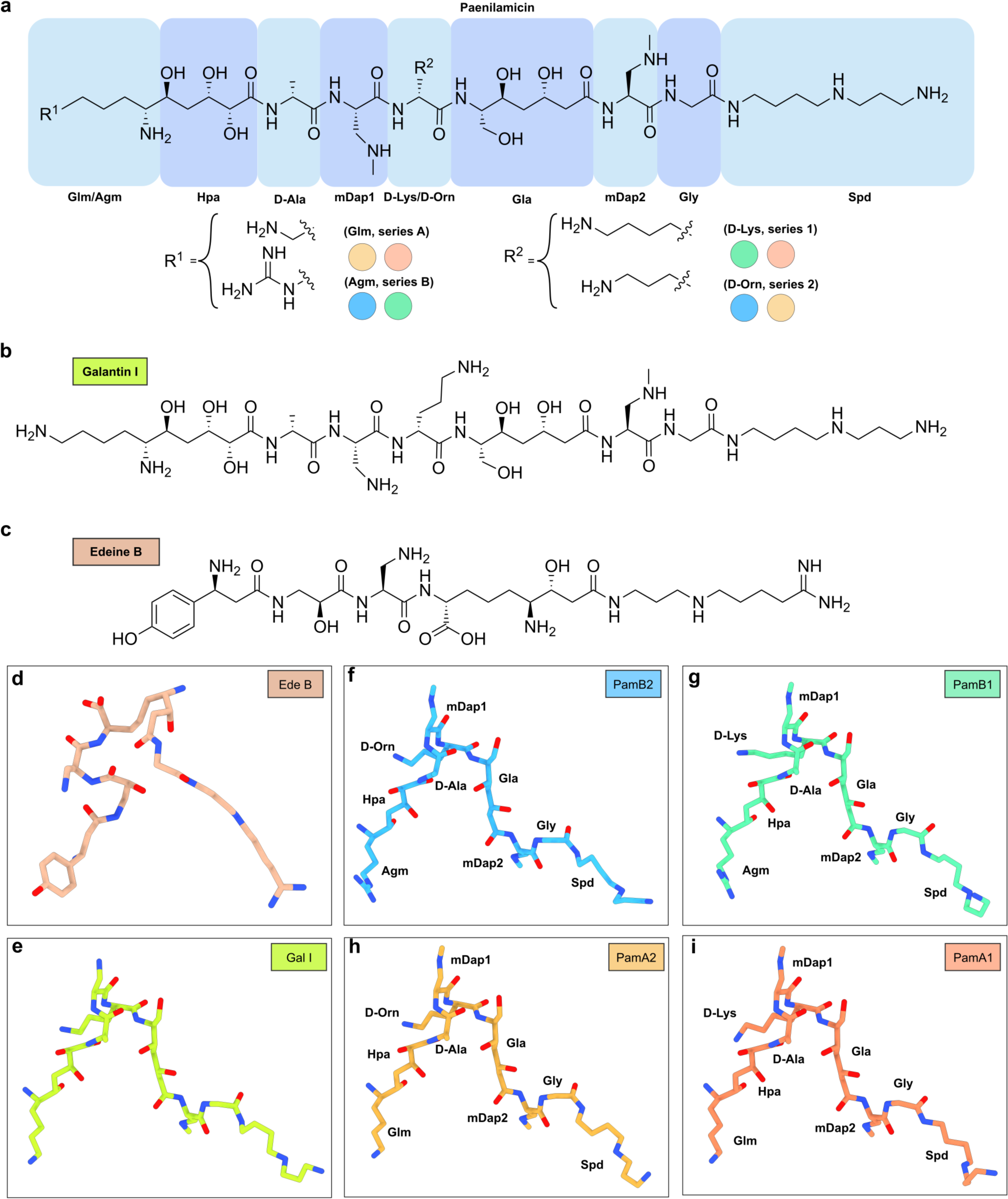
Chemical structures and models of paenilamicin, galantin I and edeine. **a-c**, Chemical structures of (**a**) paenilamicin, (**b**) galantin I and (**c**) edeine B. **d**, Molecular model of edeine B on the *T. thermophilus* 30S subunit (beige, Ede B, PDB ID 1I95)^1^. **e**, *In silico* molecular model of galantin I (Gal I, light green). **f-i**, Molecular model of PamB2 of the non-rotated PamB2 complex and *in silico* modelled PamB1 (green), PamA2 (light orange), PamA1 (orange).

**Extended Data Fig. 2:**
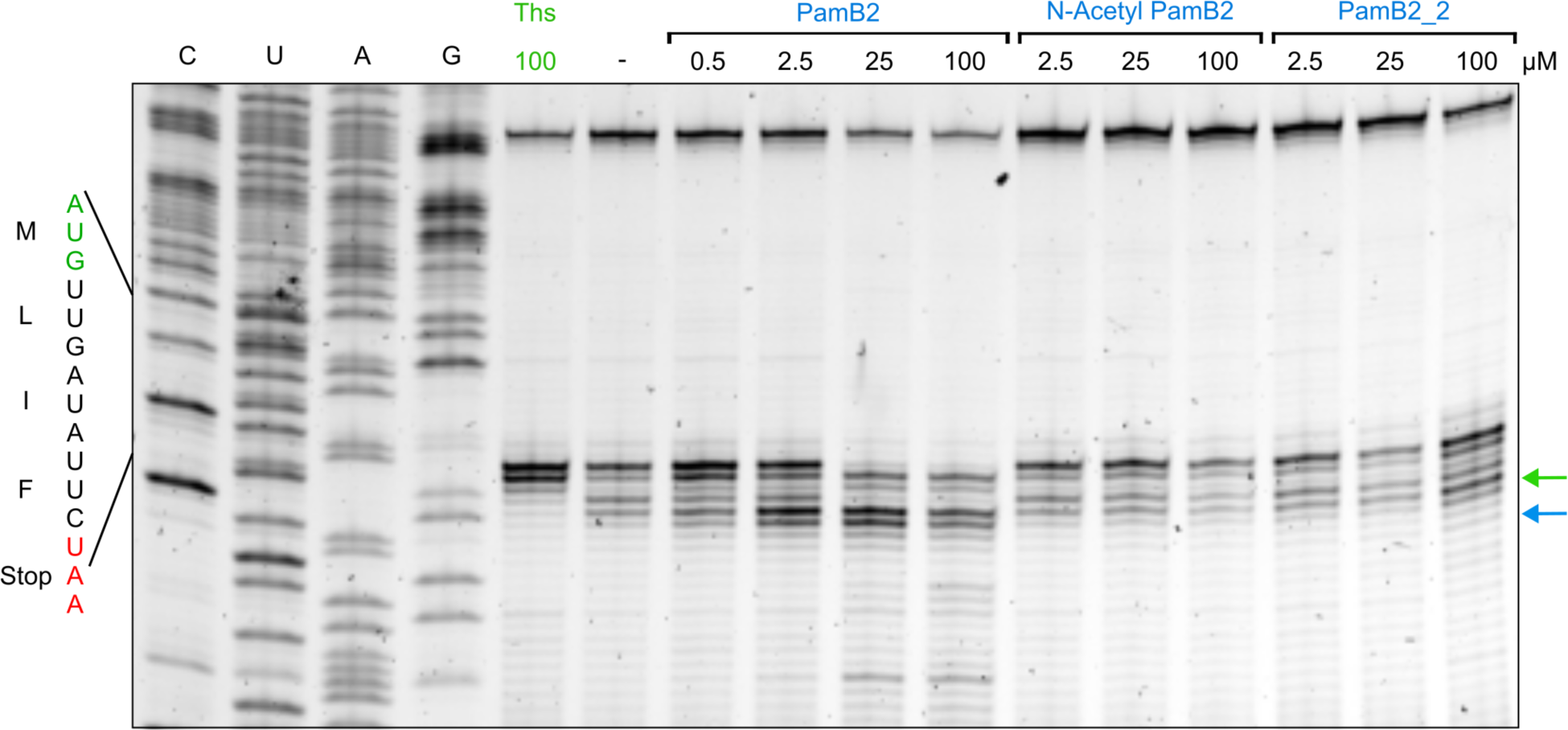
Toeprinting assay on the MLIFstop-mRNA. **a**, Toeprinting assay monitoring the position of ribosomes on a MLIFstop-mRNA in the presence of 100 µM thiostrepton, water and increasing concentrations of PamB2, N-Acetyl-PamB2 and PamB2_2 (0.5-100 µM). Arrows indicate the stalling at the initiation (green), and PamB2 induced stalling (blue). Toeprinting assays were performed in duplicate, with the duplicate gel present in the Source Data.

**Extended Data Fig. 3:**
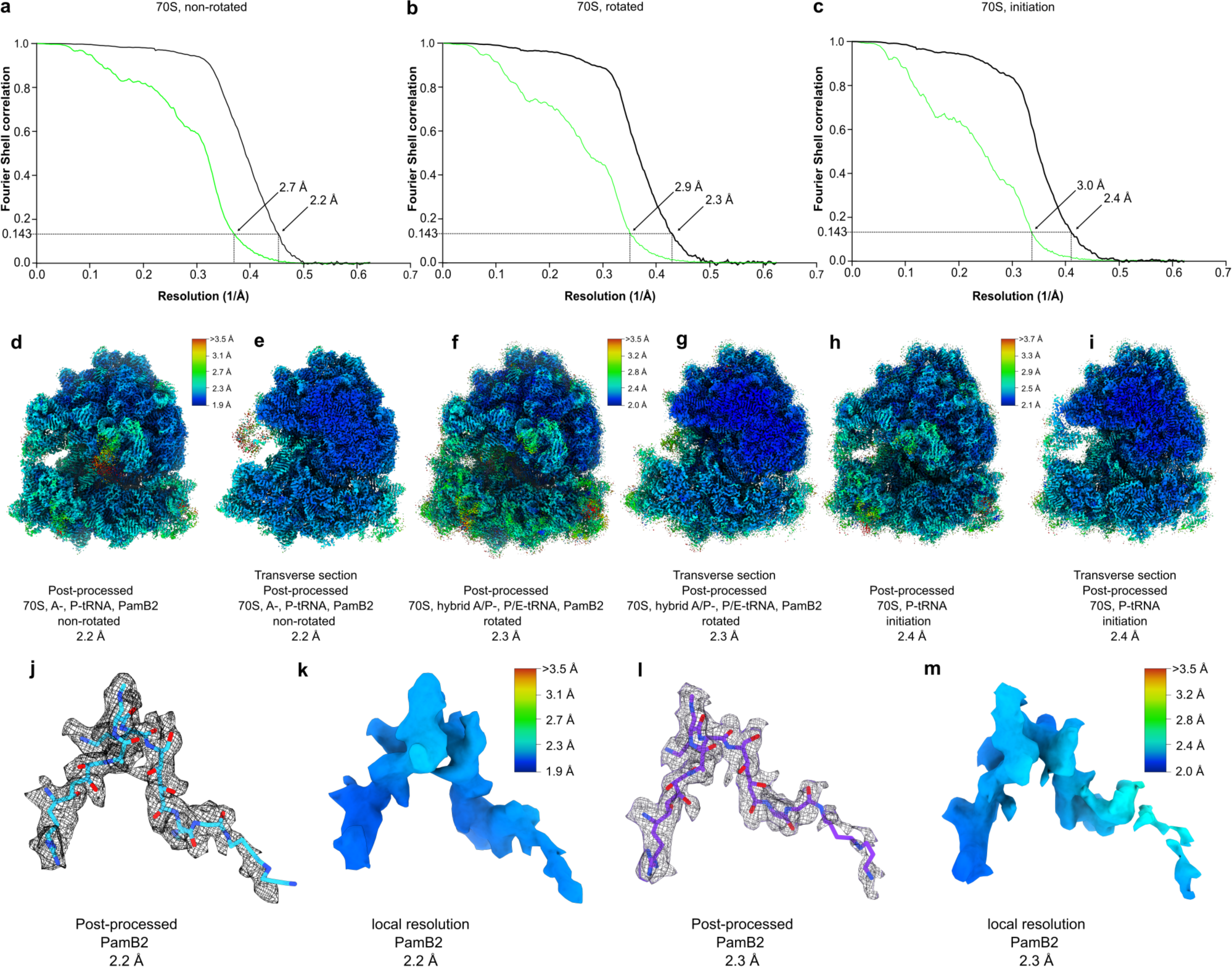
Fourier shell correlation and local resolution for the PamB2 complexes. **a-c**, Fourier shell correlation (FSC) curve of the (**a**) non-rotated, (**b**) rotated and (**c**) initiation complexes, with unmasked (green) and masked (black) FSC curves plotted against the resolution (1/Å). **d-i**, Cryo-EM density colored according to local resolution and transverse section for the (**d-e**) non-rotated, (**f-g**) rotated and (**h-i**) initiation complexes. **j-m**, Molecular model of PamB2 (light blue and purple) and corresponding cryo-EM density colored according to local resolution for the (**j-k**) non-rotated, and (**l-m**) rotated complex.

**Extended Data Fig. 4:**
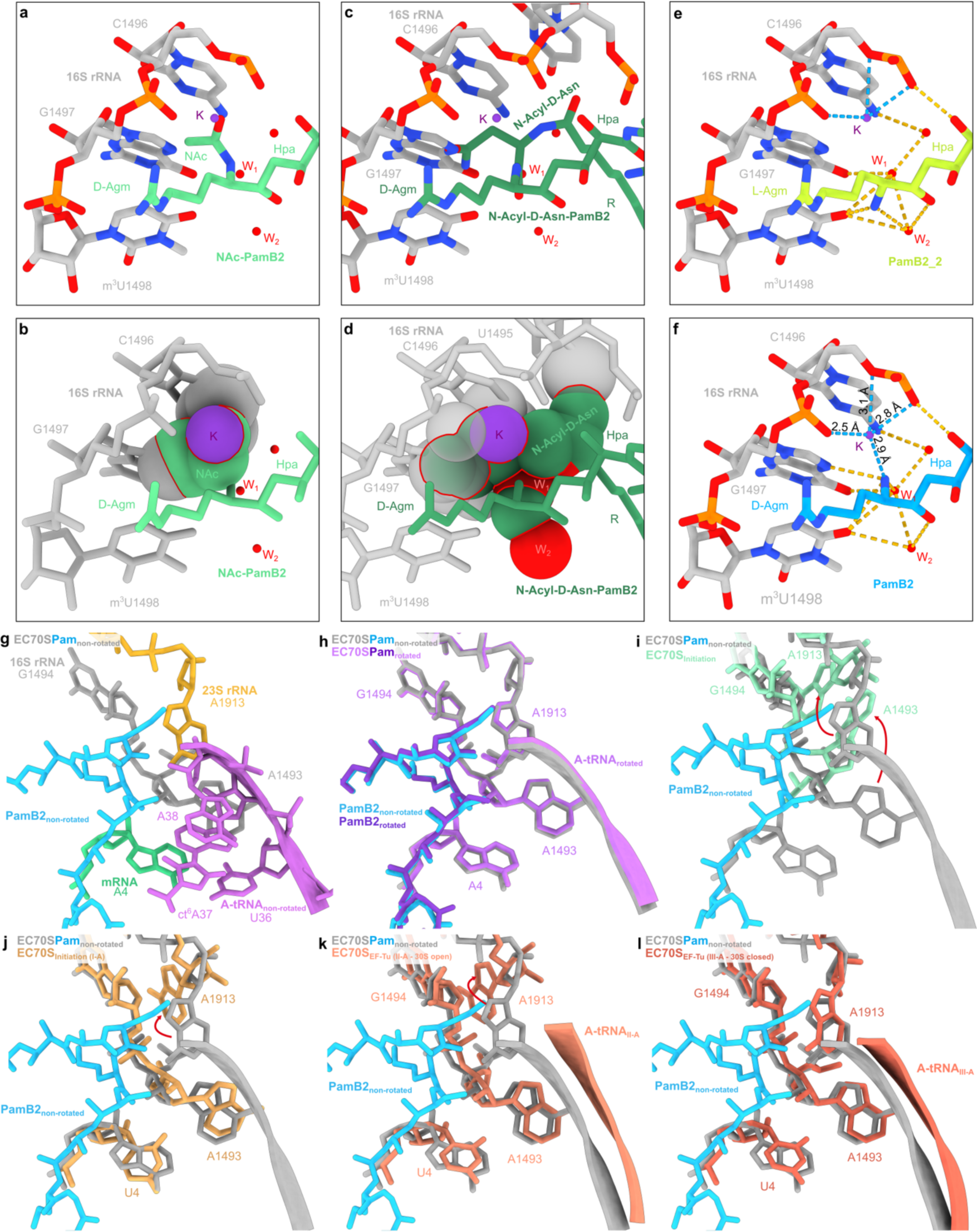
PamB2 *in silico* modification and A-site binding pocket. **a**-**d**, *In silico* model of (**a-b**) N-acetyl-PamB2 (green) shown as (**a**) stick and (**b**) sphere representation, and (**c-d**) PamB2 with an acyl-D-Asn moiety (dark green) shown as (**c**) stick and (**d**) sphere representation. In (**b**) and (**d**), the sterically clashing molecules are highlighted by red lines. **e**, Interactions of the PamB2 (light blue) wildtype D-Agm amino group with surrounding 16S rRNA of h44. **f**, Interactions of the *in silico* modified PamB2_2 (light green) L-Agm amino group with surrounding 16S rRNA of h44. **g**, Binding pocket of PamB2 (light blue) with surrounding A-site tRNA nucleotides (purple), 16S rRNA (grey), mRNA (cyan) and 23S rRNA (yellow) superimposed with (**h**) the rotated PamB2 complex (purple), (**i**) the initiation complex (light cyan),the *E. coli* initiation state (yellow, I-A, PDB ID 6WD0)^18^, (**j**) EF-Tu bound to the open *E. coli* 30S subunit (orange, II-A, PDB-ID 6WD2)^18^, and (**k**) EF-Tu bound to the closed *E. coli* 30S subunit (dark orange, III-A, PDB-ID 6WD8)^18^.

**Extended Data Fig. 5:**
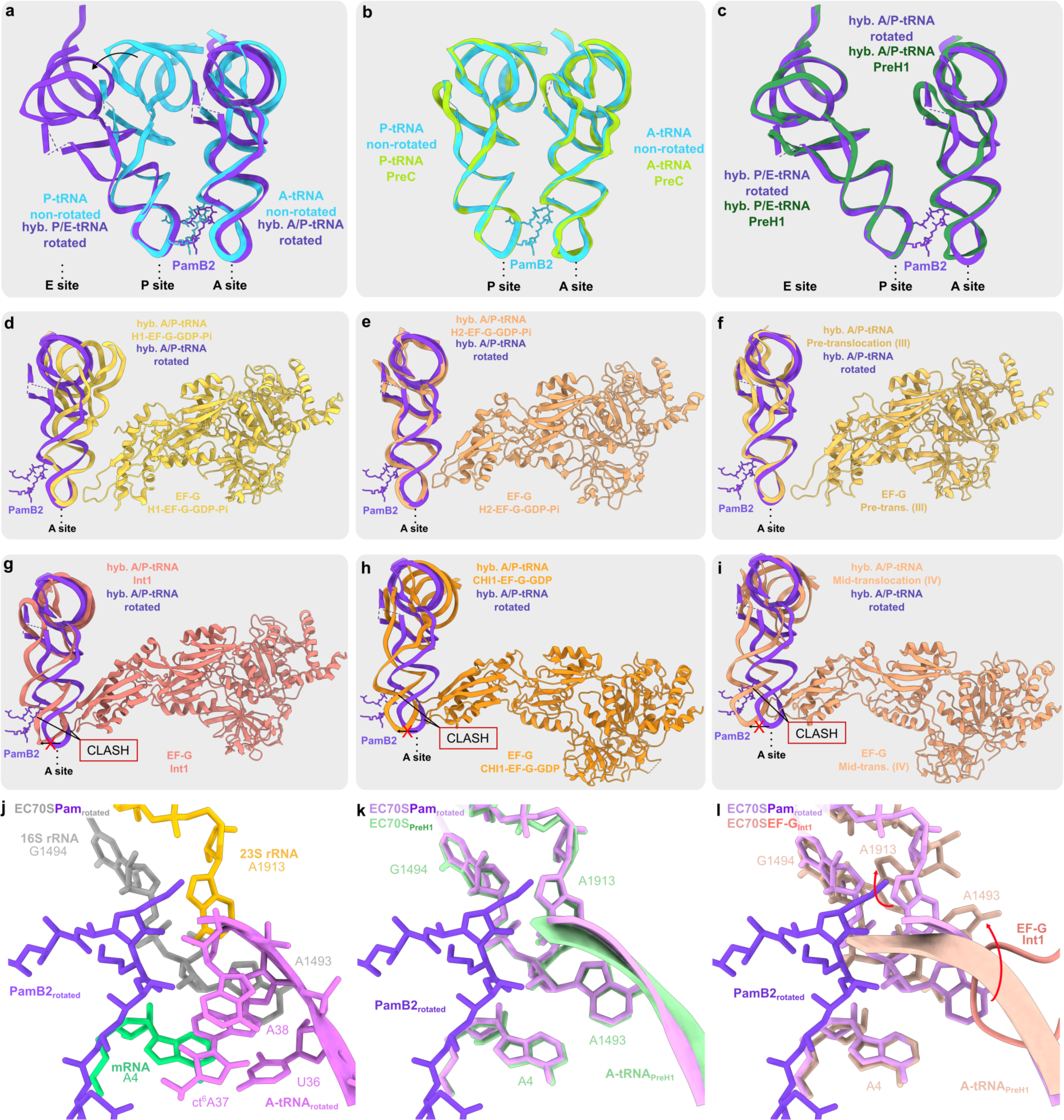
PamB2 inhibits tRNA_2_-mRNA translocation. **a**, Superimposition of the P- and A-site tRNA of the non-rotated PamB2 complex (light blue) and the hybrid A/P- and P/E-tRNA of the rotated PamB2 complex (dark purple). **b**, Superimposition of the P- and A-tRNA of the non-rotated PamB2-complex (light blue) and the P- and A-tRNA of the PreC state (light green, PDB ID 7N1P)^19^. **c**, Superimposition of the hybrid A/P- and P/E-site tRNA of the rotated PamB2-complex (dark purple) and the hybrid A/P- and P/E-site tRNA of the PreH1 state (dark green, PDB ID 7N2U)^19^. **d-i**, Superimposition of the hybrid A/P-site tRNA of the rotated PamB2-complex (dark purple) and the hybrid A/P-site tRNA and EF-G of the (**d**) H1-EF-G-GDP-Pi state (light yellow, PDB ID 7PJV)^21^, (**e**) H2-EF-G-GDP-Pi state (light orange, PDB ID 7PJW)^21^ and (**f**) pre-translocation (III) state (yellow, PDB ID 7SSL)^20^, (**g**) Int1 state (salmon, PDB ID 7N2V)^19^, (**h**) CHI1-EF-G-GDP state (orange, PDB ID 7PJY)^21^, (**i**) mid-translocation (IV) state (light orange, PDB ID 7SSD)^20^. **j-l**, PamB2 of the rotated complex (**j**) with surrounding hybrid A/P-tRNA (purple), 16S rRNA nucleotides (grey), 23S rRNA nucleotides (yellow) and the A-site codon of the mRNA (cyan) superimposed with (**k**) with the 70S *E. coli* ribosome in the PreH1 state (light green, PDB ID 7N2U)^19^ and (**l**) with the 70S *E. coli* ribosome and EF-G of the Int1 state shown as ribbon (salmon, PDB ID 7N2V)^19^.

