## Supplementary Information for "Paenilamicins from the honey bee pathogen *Paenibacillus larvae* are context-specific translocation inhibitors of protein synthesis"

**Supplementary Information**
**for**

### Paenilamicins from the honey bee pathogen *Paenibacillus larvae* are context-specific translocation inhibitors of protein synthesis

Timm O. Koller[1,#], Max J. Berger[1,#], Martino Morici[1], Helge Paternoga[1], Timur Bulatov[2], Adriana Di Stasi[3], Tam Dang[2], Andi Mainz[2], Karoline Raulf[1], Caillan Crowe-McAuliffe[1], Marco Scocchi[3], Mario Mardirossian[3], Bertrand Beckert[4], Nora Vázquez-Laslop[5], Alexander Mankin[5], Roderich D. Süssmuth[2], Daniel N. Wilson[1,*]

[1] Institute for Biochemistry and Molecular Biology, University of Hamburg, Martin-Luther-King-Platz 6, 20146 Hamburg, Germany.

[2] Institut für Chemie, Technische Universität Berlin, 10623 Berlin, Germany.

[3] Department of Life Sciences, University of Trieste, 34127 Trieste, Italy.

[4] Dubochet Center for Imaging (DCI) at EPFL, EPFL SB IPHYS DCI, Lausanne, Switzerland.

[5] Center for Biomolecular Sciences, University of Illinois at Chicago, Chicago, IL 60607.

[#] These authors contributed equally.

*Correspondence to:

Daniel Wilson

**Contents include:**

Supplementary Figures 1-5
Supplementary Tables 1-3
Supplementary References

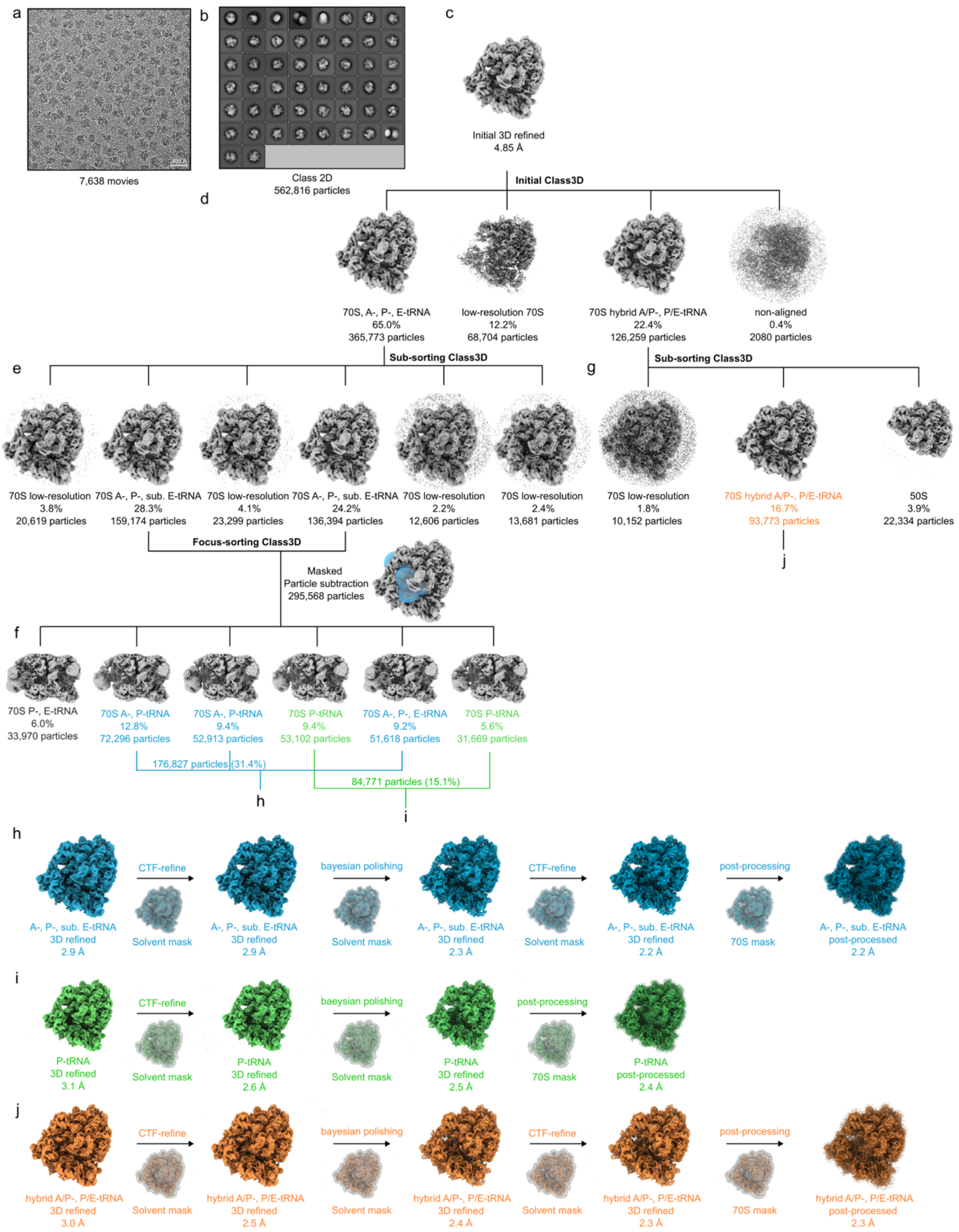

**Supplementary Fig. 1: *In silico* sorting scheme of the *E. coli* 70S PamB2 complex. a-b**, From 7,638 micrographs (**a**) a total of 562,816 ribosome-like particles were selected after 2D classification (**b**). **c**, Particles were subjected to an initial 3D refinement at 3x decimated pixel size. **d**, Particles were 3D classified for 100 iterations and resulted in four classes of which a non-rotated 70S class with A-, P- and E-site tRNAs (65.0%, 365,773 particles) and a rotated 70S with hybrid A/P- and P/E-tRNA (22.4%, 126,259 particles) were further sub-sorted. **e**, The non-rotated 70S class was 3D classified for 100 iterations and yielded six classes of which four classes were low resolution 70S particles and two classes of 70S with A-, P- and sub-stoichiometric E-site tRNA (52.5%, 295,568 particles) were combined. **f**, The combined classes from (**e**) were partially subtracted with a mask surrounding the tRNAs and 3D classified for 100 iterations yielding six classes. Classes containing A-, P- and E-site tRNA (blue, 31.4%, 176,827 particles), as well as classes containing just P-site tRNA (green, 15.1%, 84,771 particles) were combined and further processed. **g**, The rotated 70S class was 3D classified for 100 iterations and yielded three classes which contained rotated 70S with hybrid A/P- and P/E-tRNA (orange, 16.7%, 93,773 particles), just 50S subunits (3.9%, 22,334 particles) and low resolution particles. The class containing rotated 70S was further processed. **h-j**, Particles were 3D refined at undecimated pixel size with a solvent mask, subjected to CTF refinement ($4^{th}$ order aberrations, anisotropic magnification and per-particle defocus value estimation), Bayesian polished, again CTF refined and after a final 3D refinement yielded (**h**) a final average resolution of 2.2 Å (at $FSC_{0.143}$) for the post-processed masked reconstruction of the non-rotated 70S complex containing A-, P- and sub. E-site tRNAs (blue), (**i**) a final average resolution of 2.4 Å (at $FSC_{0.143}$) for the post-processed masked reconstruction of the 70S complex containing P-tRNA (green) and (**j**) a final average resolution of 2.3 Å (at $FSC_{0.143}$) for the post-processed masked reconstruction of the rotated 70S complex containing hybrid A/P-, and P/E-tRNAs (orange). The single particles analysis was performed in RELION and a single time for 3D refinements, and employing the gold-standard, particles are randomly placed in one of two subsets and processed independently for both half-reconstructions. These subsets are maintained for CTF refinement.

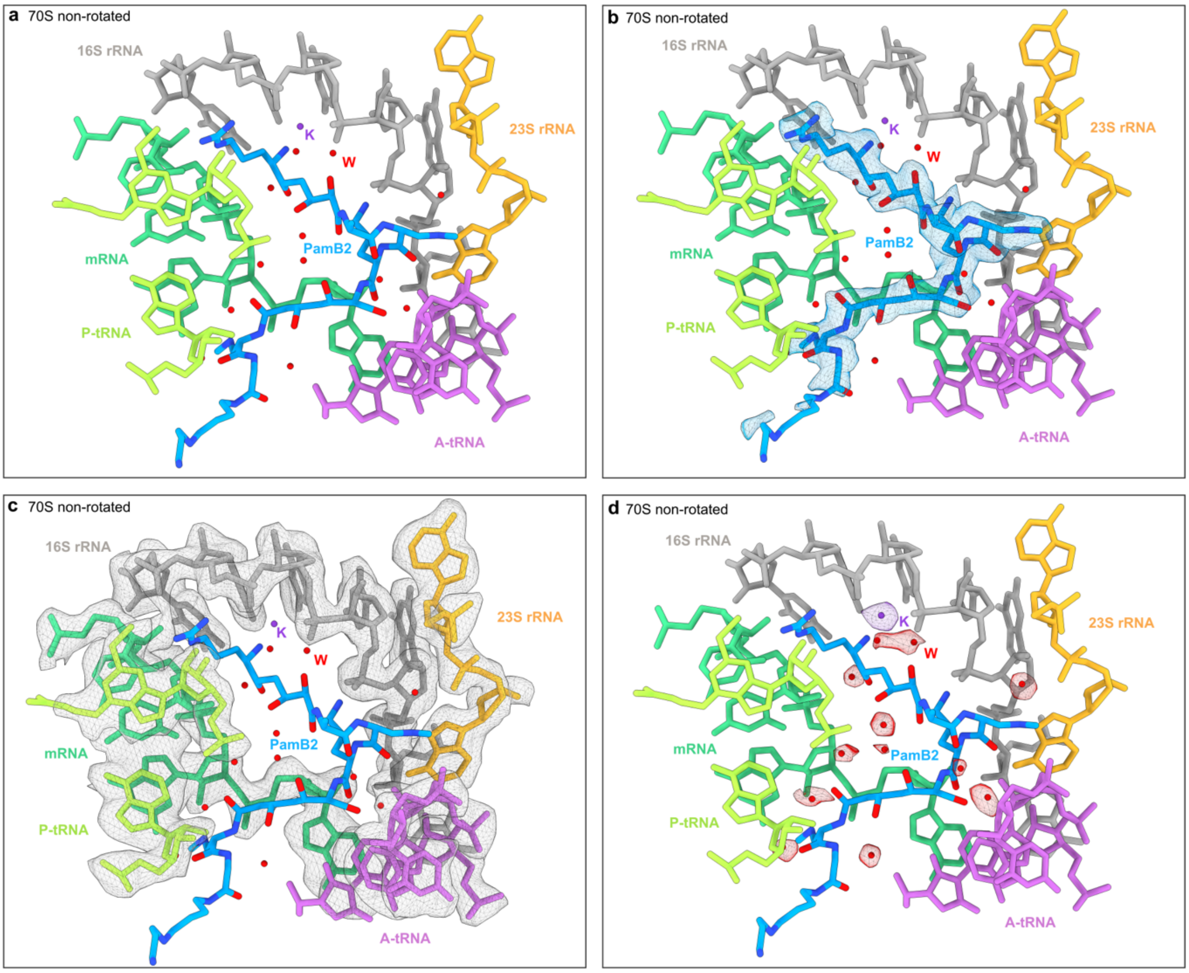

**Supplementary Fig. 2: Cryo-EM density of the PamB2 binding pocket. a-d**, PamB2 (blue) binding pocket surrounded by 16S rRNA nucleotides (grey), 23S rRNA nucleotides (yellow), mRNA (cyan), A- (purple) and P-site tRNA (light green), and waters (red) and a potassium ion (dark purple). Cryo-EM density of the non-rotated 70S PamB2 complex is shown as mesh for extracted density at one threshold for (**b**) PamB2, (**c**) surrounding nucleotides and (**d**) waters and potassium ion.

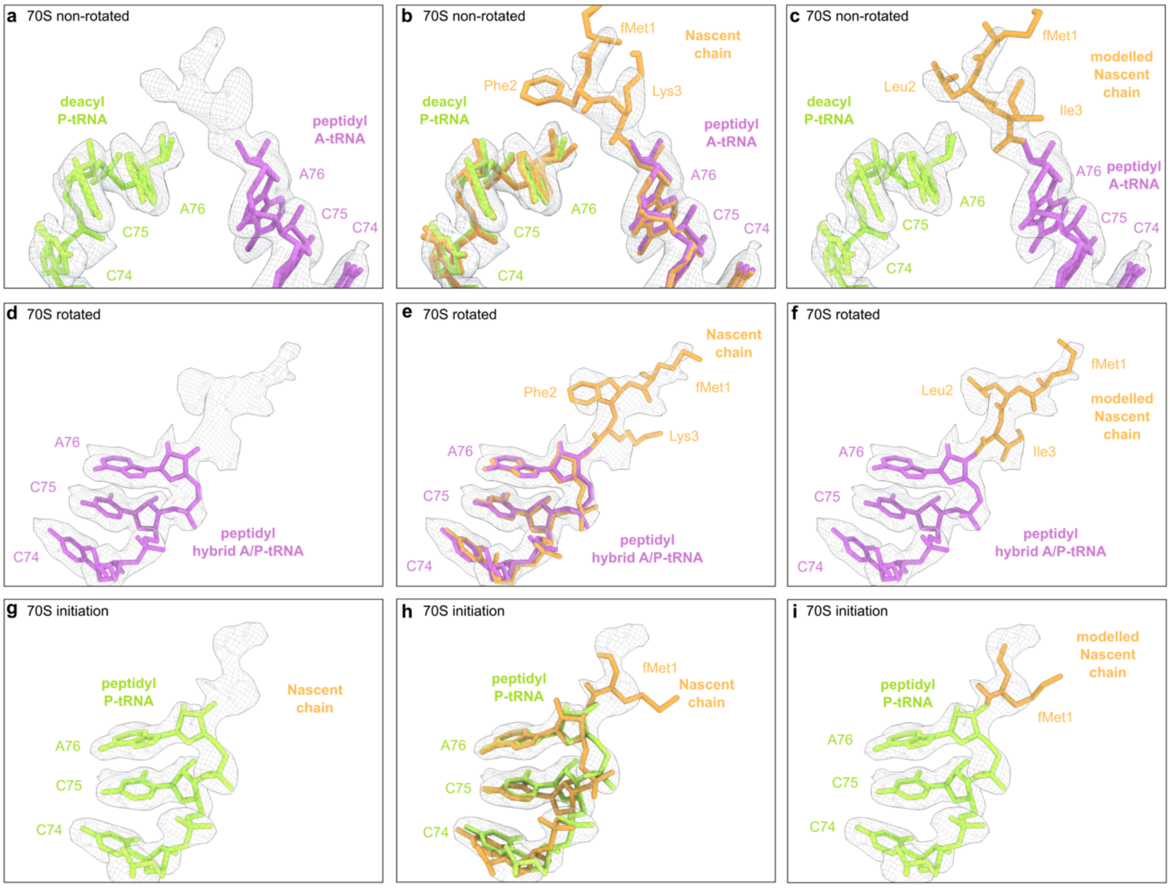

**Supplementary Fig. 3: Additional cryo-EM density for the nascent chain attached to the CCA-end of the peptidyl-tRNA. a-i**, Extracted cryo-EM densities of the respective complexes were shown as mesh with P-site tRNA (light green), A-site tRNA (purple) and hybrid A/P-site tRNA (purple). **a-c**, Additional cryo-EM density (**a**) on the A-site tRNA was superimposed with (**b**) a tri-peptide of a non-rotated 70S *E. coli* nascent chain from the PreC state (orange, PDB ID 7N1P)[1] and (**c**) a hypothetical molecular model of the fMet-Leu-Ile nascent chain connected to the A-site tRNA. **d-f**, Additional cryo-EM density (**d**) on the hybrid A/P-site tRNA was superimposed with (**e**) a tri-peptide of a rotated 70S *E. coli* nascent chain from the PreH1 state (orange, PDB ID 7N2U)[1] and (**f**) a hypothetical molecular model of the fMet-Leu-Ile nascent chain connected to the hybrid A/P-site tRNA. **g-h**, Additional cryo-EM density (**d**) on the P-site tRNA of the 70S initiation complex was superimposed with (**h**) an initiator fMet-tRNA$^{fMet}$ from *an E. coli* 70S initiation state (orange, PDB ID 6WD0)[2] and (**f**) a hypothetical molecular model of the fMet nascent chain connected to the P-site tRNA.

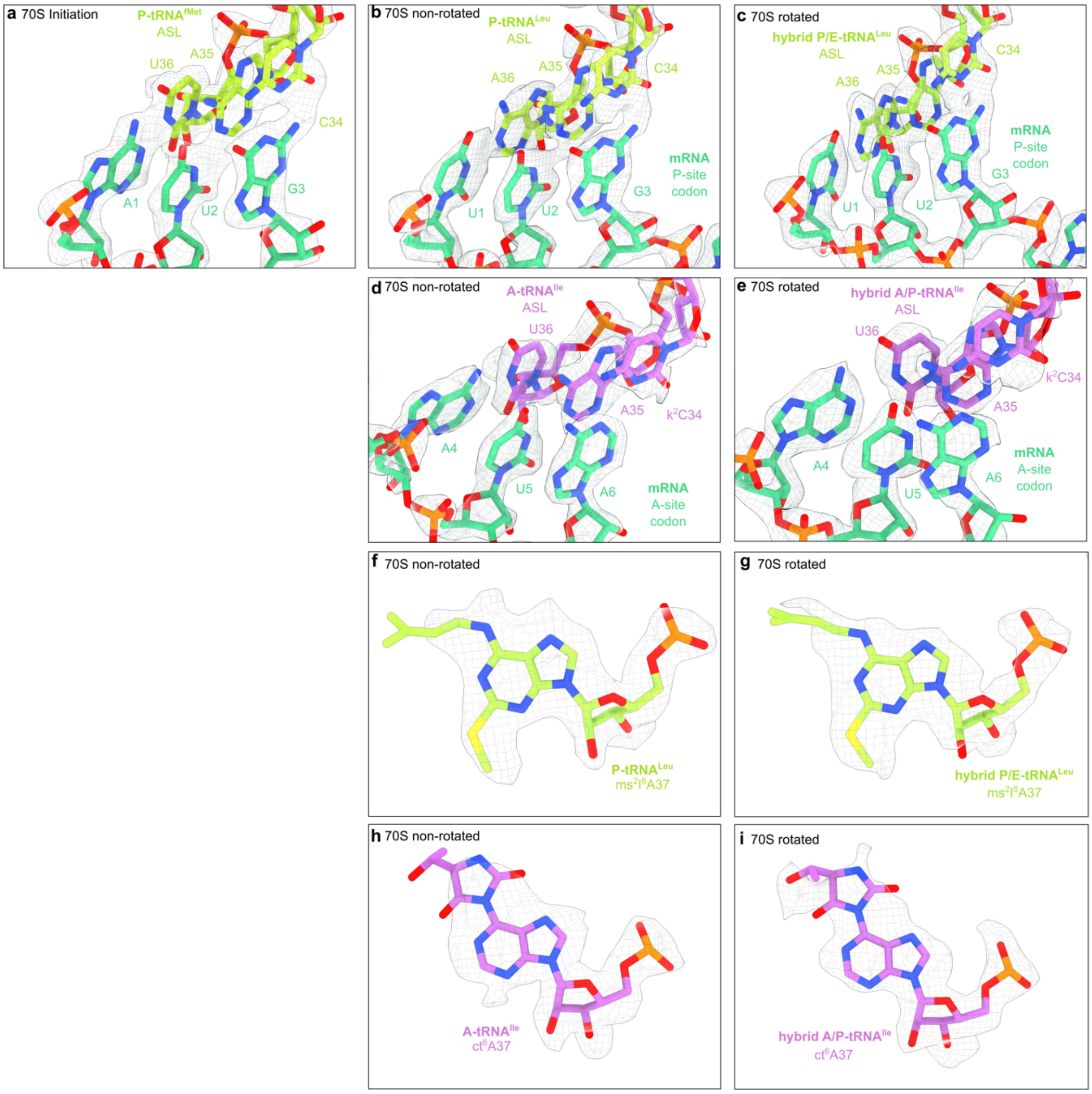

**Supplementary Fig. 4: Codon-anticodon-interaction and tRNA identity. a-c**, Extracted density of the codon-anticodon-interaction of the (**a**) P-site tRNA$^{fMet}$ anticodon-stem loop (light green) and P-site codon (cyan, AUG) of 70S initiation complex, (**b**) P-site tRNA$^{Leu}$ anticodon-stem loop (light green) and P-site codon (cyan, UUG) of the 70S non-rotated PamB2 complex and (**c**) hybrid P/E-tRNA$^{Leu}$ anticodon-stem loop (light green) and P-site codon (cyan, UUG) of the 70S rotated PamB2 complex. **d-e**, Extracted density of the codon-anticodon-interaction of the (**d**) A-site tRNA$^{Ile}$ anticodon-stem loop (purple) and A-site codon (cyan, AUA) of the 70S non-rotated complex and (**e**) hybrid A/P-tRNA$^{Ile}$ anticodon-stem loop (purple) and P-site codon (cyan, UUG) of the 70S rotated PamB2 complex. **f-g**, Extracted cryo-EM density of the 2-methylhio-N6-isopentenyladenine (ms$^2$i$^6$, light green) modification at position 37 of the (**f**) P-site tRNA$^{Leu}$ of the non-rotated PamB2 complex and the (**g**) hybrid P/E-site tRNA of the rotated PamB2 complex. **h-i**, Extracted cryo-EM density of the cyclic N6-threonylcarbamoyladenine (ct6, purple) modification at position 37 of the (**h**) A-site tRNA$^{Ile}$ of the non-rotated PamB2 complex and the (**i**) hybrid A/P-site tRNA$^{Ile}$ of the rotated PamB2 complex.

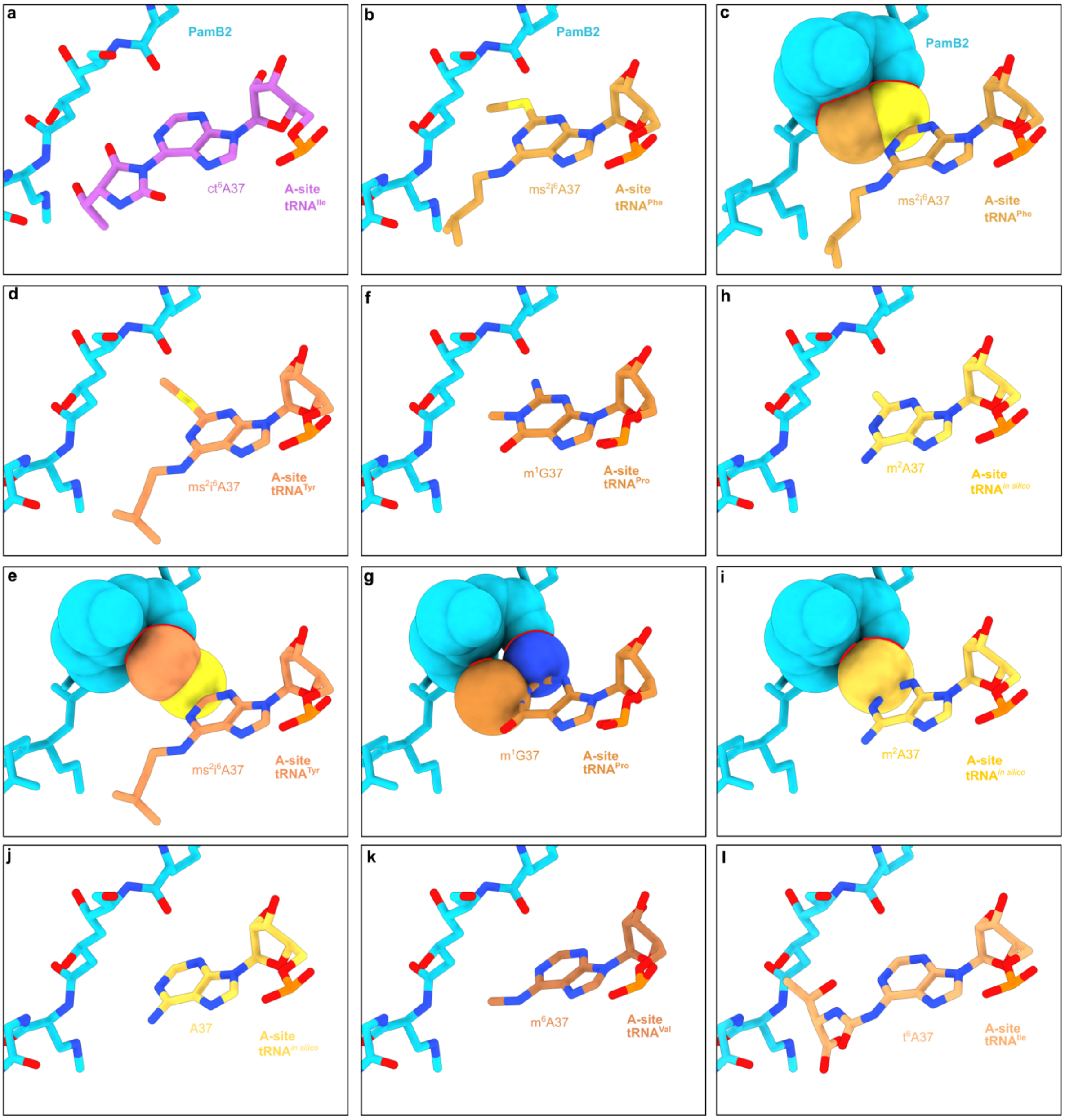

**Supplementary Fig. 5: Influence of A37 modification of A-tRNA on PamB2 inhibition. a-l**, PamB2 (light blue) from the non-rotated PamB2 complex superimposed with tRNAs with modified residues at position 37. **a**, The cyclic N6-threonylcarbamoyladenine (ct6, purple) modification at position 37 of the A-site $tRNA^{Ile}$ from the non-rotated PamB2 complex. **b-c**, PamB2 from (**a**) superimposed with (**b**) $tRNA^{Phe}$ with a 2-methylhio-N6-isopentenyladenine ($ms^2i^6$, light orange, PDB ID 1VY5)[3] at position 37 and sphere representation with (**c**) steric clashes highlighted in red. **d-e**, PamB2 from (**a**) superimposed with (**d**) $tRNA^{Tyr}$ with a 2-methylhio-N6-isopentenyladenine ($ms^2i^6$, orange, PDB ID 4V8D)[4] at position 37 and sphere representation with (**e**) steric clashes highlighted in red. **f-g**, PamB2 from (**a**) superimposed with (**b**) $tRNA^{Tyr}$ with a 1-methyl-guanine ($m^1G$, dark orange, PDB ID 6NUO)[5] at position 37 and sphere representation with (**g**) steric clashes highlighted in red. **h-i**, PamB2 from (**a**) superimposed with (**h**) an *in silico* modified tRNA with 2-methyl-adenine ($m^2A$, yellow) at position 37 and sphere representation with (**i**) steric clashes highlighted in red. **j,** PamB2 from (**a**) superimposed with an *in silico* tRNA with unmodified A37. **k**, PamB2 from (**a**) superimposed with $tRNA^{Val}$ with 6-methyl-adenine ($m^6A$, brown, PDB ID 4V6Z)[6] in position 37. **i**, PamB2 from (**a**) superimposed with $tRNA^{Ile}$ with N6-threonylcarbamyl ($t^6A$, beige, PDB ID 7N1P)[1] in position 37.

**Supplementary Table 1. Cryo-EM data collection, modelling and refinement statistics.**

| | **Non-rotated 70S PamB2 complex** (EMD-18950) (PDB ID 8R6C) | **Rotated 70S PamB2 complex** (EMD-19004) (PDB ID 8R8M) | **70S Initiation complex** (EMD-50296) (PDB ID 9FBV) |
|---|---|---|---|
| **Data collection** | | | |
| Magnification (×) | 96,000 | 96,000 | 96,000 |
| Voltage (kV) | 300 | 300 | 300 |
| Electron exposure ($e^{-}$/Å$^2$) | 60 | 60 | 60 |
| Defocus range (μm) | -0.4 to -0.9 | -0.4 to -0.9 | -0.4 to -0.9 |
| Pixel size (Å) | 0.80 | 0.80 | 0.80 |
| Symmetry imposed | C1 | C1 | C1 |
| Initial particle images (no.) | 562,816 | 562,816 | 562,816 |
| Final particle images (no.) | 176,827 | 93,773 | 84,771 |
| Map resolution (Å) | 2.2 | 2.3 | 2.4 |
| FSC threshold | 0.143 | 0.143 | 0.143 |
| Map resolution range (Å) | 1.9-3.2 | 2.0-3.5 | 2.1-4.0 |
| **Refinement** | | | |
| Initial model used (PDB) | 7K00 | 7K00 | 7K00 |
| Model resolution (Å) | 2.6 | 2.7 | 2.9 |
| FSC threshold | 0.5 | 0.5 | 0.5 |
| Model resolution range (Å) | 2.0-3.2 | 2.1-3.5 | 2.2-3.5 |
| Map sharpening B factor (Å$^2$) | -4.74 | -7.43 | -8.04 |
| Model composition | | | |
| Non-hydrogen atoms | 145,011 | 141,381 | 140,097 |
| Protein residues | 5,585 | 5,573 | 5,585 |
| RNA bases | 4,547 | 4,530 | 4,470 |
| *B* factors (Å$^2$) | | | |
| Protein | 74.9 | 76.2 | 66.4 |
| Nucleotide | 64.7 | 62.1 | 54.8 |
| R.M.S. deviations | | | |
| Bond lengths (Å) | 0.010 | 0.010 | 0.010 |
| Bond angles (°) | 1.430 | 1.484 | 1.418 |
| Validation | | | |
| MolProbity score | 1.17 | 1.35 | 1.64 |
| Clash score | 0.63 | 0.90 | 1.92 |
| Poor rotamers (%) | 2.10 | 2.87 | 3.70 |
| Ramachandran statistics | | | |
| Favoured (%) | 96.64 | 96.49 | 96.20 |
| Allowed (%) | 3.05 | 3.15 | 3.58 |
| Disallowed (%) | 0.31 | 0.37 | 0.20 |

**Supplementary Table 2. Primers used in this study.**

| Primer Name | Sequence (5’ -3’) |
|---|---|
| **ErmBL-UGA-NV1-R** | GGTTATAATGAATTTTGCTTATTAACGATAGAATTCTATC ACTCAAATAGTAGATGTTTTATCTACATTACGCATTT |
| **T7-F** | TAATACGACTCACTATAGGG |
| **T7-ErmBL- F** | TAATACGACTCACTATAGGGAGACTTAAGTATAAGGAGG AAAAAATATGTTGGTATTCCAAATGCGTAATGTAGATAA |
| **UUG1 -F** | TTAGTATAAGGAGGAAAAAATATGTTGGTATTCCAAATGC GTAATG |
| **UUG2 -F** | TTAGTATAAGGAGGAAAAAATATGTTGTTGGTATTCCAAAT GCGTAATG |
| **UUG3 -F** | TTAGTATAAGGAGGAAAAAATATGTTGTTGTTGGTATTCCA AATGCGTAATG |
| **UUG4 -F** | TTAGTATAAGGAGGAAAAAATATGTTGTTGTTGTTGGTATT CCAAATGCGTAATG |
| **UUG5 -F** | TTAGTATAAGGAGGAAAAAATATGTTGTTGTTGTTGTTGGT ATTCCAAATGCGTAATG |
| **T7-SD-AUG-F** | TAATACGACTCACTATAGGGCTTAGTATAAGGAGGAAAAAA TATG |
| **UUG4-UCU-F** | TTAGTATAAGGAGGAAAAAATATGTTGTTGTTGTTGTCTTT CCAAATGCGTAATGTAG |
| **UUG4-UCC-F** | TTAGTATAAGGAGGAAAAAATATGTTGTTGTTGTTGTCCTT CCAAATGCGTAATGTAG |
| **UUG4-UCA-F** | TTAGTATAAGGAGGAAAAAATATGTTGTTGTTGTTGTCATT CCAAATGCGTAATGTAG |
| **UUG4-UCG-F** | TTAGTATAAGGAGGAAAAAATATGTTGTTGTTGTTGTCGTT CCAAATGCGTAATGTAG |
| **NV1-ErmBL-R** | GGTTATAATGAATTTTGCTTATTAACGATAGAATTCTATCAC TTACAAAATAGTAGATGTGATTTTATCTACATTACGCATTTG GAATAC |
| **NV1** | GGTTATAATGAATTTTGCTTATTAACC |

**Supplementary Table 3. mRNA templates used in this study.**

| Template Name | Sequence (5’ -3’) |
|---|---|
| **ErmBL** | TAATACGACTCACTATAGGGAGACTTAAGTATAAGGAGGAAAAA<br>AT**ATG**TTGGTATTCCAAATGCGTAATGTAGATAAAACATCTACTA<br>TTTGAGTGATAGAATTCTATCGTTAATAAGCAAAATTCATTATAAC<br>C |
| **(UUG)2-ErmBL** | TAATACGACTCACTATAGGGCTTAGTATAAGGAGGAAAAAAT**AT**<br>**G**TTGTTGGTATTCCAAATGCGTAATGTAGATAAAATCACATCTAC<br>TATTTTGTAAGTGATAGAATTCTATCGTTAATAAGCAAAATTCATT<br>ATAACC |
| **(UUG)$_3$-ErmBL** | TAATACGACTCACTATAGGGCTTAGTATAAGGAGGAAAAAAT**AT**<br>**G**TTGTTGTTGGTATTCCAAATGCGTAATGTAGATAAAATCACATC<br>TACTATTTTGTAAGTGATAGAATTCTATCGTTAATAAGCAAAATTC<br>ATTATAACC |
| **(UUG)$_4$-ErmBL** | TAATACGACTCACTATAGGGCTTAGTATAAGGAGGAAAAAAT**AT**<br>**G**TTGTTGTTGTTGGTATTCCAAATGCGTAATGTAGATAAAATCAC<br>ATCTACTATTTTGTAAGTGATAGAATTCTATCGTTAATAAGCAAAA<br>TTCATTATAACC |
| **(UUG)$_5$-ErmBL** | TAATACGACTCACTATAGGGCTTAGTATAAGGAGGAAAAAAT**AT**<br>**G**TTGTTGTTGTTGTTGGTATTCCAAATGCGTAATGTAGATAAAAT<br>CACATCTACTATTTTGTAAGTGATAGAATTCTATCGTTAATAAGCA<br>AAATTCATTATAACC |
| **(UUG)$_4$-UCU-ErmBL** | TAATACGACTCACTATAGGGCTTAGTATAAGGAGGAAAAAAT**AT**<br>**G**TTGTTGTTGTTGTCTTTCCAAATGCGTAATGTAGATAAAATCAC<br>ATCTACTATTTTGTAAGTGATAGAATTCTATCGTTAATAAGCAAAA<br>TTCATTATAACC |
| **(UUG)$_4$-UCC-ErmBL** | TAATACGACTCACTATAGGGCTTAGTATAAGGAGGAAAAAAT**AT**<br>**G**TTGTTGTTGTTGTCCTTCCAAATGCGTAATGTAGATAAAATCAC<br>ATCTACTATTTTGTAAGTGATAGAATTCTATCGTTAATAAGCAAAA<br>TTCATTATAACC |
| **(UUG)$_4$-UCA-ErmBL** | TAATACGACTCACTATAGGGCTTAGTATAAGGAGGAAAAAAT**AT**<br>**G**TTGTTGTTGTTGTCATTCCAAATGCGTAATGTAGATAAAATCAC<br>ATCTACTATTTTGTAAGTGATAGAATTCTATCGTTAATAAGCAAAA<br>TTCATTATAACC |
| **(UUG)$_4$-UCG-ErmBL** | TAATACGACTCACTATAGGGCTTAGTATAAGGAGGAAAAAAT**AT**<br>**G**TTGTTGTTGTTGTCGTTCCAAATGCGTAATGTAGATAAAATCAC<br>ATCTACTATTTTGTAAGTGATAGAATTCTATCGTTAATAAGCAAAA<br>TTCATTATAACC |
| **ErmBL-ATG** | TAATACGACTCACTATAGGGAGACTTAAGTATAAGGAGGAAAAA<br>AT**ATG**ATGGTATTCCAAATGCGTAATGTAGATAAAACATCTACTA<br>TTTGAGTGATAGAATTCTATCGTTAATAAGCAAAATTCATTATAAC<br>C |
| **ErmBL-CTG** | TAATACGACTCACTATAGGGAGACTTAAGTATAAGGAGGAAAAA<br>AT**ATG**CTGGTATTCCAAATGCGTAATGTAGATAAAACATCTACTA<br>TTTGAGTGATAGAATTCTATCGTTAATAAGCAAAATTCATTATAAC<br>C |
| **ErmBL-GTG** | TAATACGACTCACTATAGGGAGACTTAAGTATAAGGAGGAAAAA<br>AT**ATG**GTGGTATTCCAAATGCGTAATGTAGATAAAACATCTACTA<br>TTTGAGTGATAGAATTCTATCGTTAATAAGCAAAATTCATTATAAC<br>C |
| **MLIF-UAA** | TAATACGACTCACTATAGGGAGACTTAAGTATAAGGAGGAAAAA<br>AT**ATG**TTGATATTCTAAATGCGTAATGTAGATAAAACATCTACTAT<br>TTAAGTGATAGAATTCTATCGTTAATAAGCAAAATTCATTATAACC |

| | |
|---|---|
| **MFKAFK** | ATTAATACGACTCACTATAGGGCAACCTAAAACTTACACACGCCCCGGTAAGGAAATAAAA**ATG**TTCAAAGCATTCAAAAACATCATACGTACTCGTACTCTTTAAGCGCAGGCAAGGTTAATAAGCAAAATTCATTATAACC |
